# Population genomics reveal distinct and diverging populations of *An. minimus* in Cambodia – a widespread malaria vector in Southeast Asia

**DOI:** 10.1101/2021.11.11.468219

**Authors:** Brandyce St. Laurent, Nick Harding, Nick Deason, Kolthida Oy, Chea Sok Loeun, Men Sary, Rous Sunly, Sen Nhep, Eleanor Drury, Kirk Rockett, Siv Sovannaroth, Sonia Goncalves, Dominic Kwiatkowski, Alistair Miles

## Abstract

*Anopheles minimus* is an important malaria vector throughout its wide geographic range across Southeast Asia. Genome sequencing could provide important insights into the unique malaria transmission dynamics in this region, where many vector species feed and rest outdoors. We describe results from a study using Illumina deep whole-genome sequencing of 302 wild-caught *An. minimus* collected from three Cambodian provinces over several years (2010, 2014-2016) and seasons to examine the level of population structure and genetic diversity within this species. These specimens cluster into four distinct populations of *An. minimus s.s.*, with two populations overlapping geographically. We describe the underlying genetic diversity and divergence of these populations and investigated the genetic variation in genes known to be involved in insecticide resistance. We found strong signals of selection within these *An. minimus* populations, most of which were present in the two Northeastern Cambodian populations and differ from those previously described in African malaria vectors. Cambodia is the focus of the emergence and spread of drug-resistant malaria parasites, so understanding the underlying genetic diversity and resilience of the vectors of these parasites is key to implementing effective malaria control and elimination strategies. These data are publicly available as part of the MalariaGEN Vector Observatory, an open access resource of genome sequence data.

## Introduction

*An. minimus* is a small mosquito that plays a major role in sustaining malaria transmission across the Greater Mekong Subregion (GMS). This species is commonly associated with human-altered habitats, able to breed in rice paddies and forest-fringe habitats. The GMS is becoming increasingly fragmented as deforestation and agriculture create patchwork landscapes (*1–3*). Historically, malaria transmission in this region has been dominated by the major vector species *An. dirus*, a forest-dwelling, highly competent vector and human-attracted mosquito. More recently, land use change has resulted in the decline of *An. dirus* populations and has shed light on the role of other vectors in malaria transmission, species like *An. minimus*, that are able to breed in human-associated habitats and quickly adapt to changing environments (*4–6*). Vectors that occupy different seasonal and ecological niches can also sustain malaria transmission and change the dynamics of seasonal transmission patterns. While *An. dirus* is considered the major vector in the greater Mekong Subregion (GMS), there are many other species that have been shown to be susceptible to malaria parasites that play a role in local transmission (*2, 7–9*).

To further complicate malaria transmission in this region, successful lineages of multi-drug resistant malaria parasites have continued to emerge from the area along the Thai-Cambodian border (*10–13*) and spread through the GMS. Lab infection experiments have shown that *Anopheles minimus* is able to be infected by and transmit the parasites in Cambodia that have been shown to have slow clearance in response to the Artemisinin combined therapy (ACT) drug treatment (*14*) and these parasites are co-circulating in areas where *An. minimus* is active and abundant. All five human malarias are transmitted in Cambodia (*15*), with *Plasmodium vivax* infections about as prevalent as those from *Plasmodium falciparum* (*16*). *An. minimus* is known to transmit *P. falciparum* and *P. vivax* throughout its wide geographic range, which spans from China down into the Malay archipelago, and from western India to Vietnam (*17*).

Many *Anopheles* species in Southeast Asia are members of cryptic species complexes, groups of isomorphic species that are molecularly and often behaviourally distinct. In other *Anopheles* cryptic species groups like the Gambiae complex, some closely related cryptic species have been shown to exchange genetic material from almost every region of the genome, described as having “porous species boundaries” (*18*). The level of gene flow between what is typically thought of as isolated species has important implications for the spread of genes relevant to malaria transmission, such as insecticide resistance alleles, or genes that confer susceptibility or resistance to malaria parasites. Population genomic studies of *An. gambiae s.l*. across the African continent have shown that geographically distinct populations have similar signatures of selection in response to decades of insecticide-based vector control (*19*), sharing common haplotypes that confer insecticide resistance over thousands of kilometers. *Anopheles minimus* is a member of such a species complex with at least three recognized cryptic species: *An. minimus* s.s. (*An. minimus* A), *An. harrisoni (An. minimus C*), and *An. yaeyamaensis* (*An. minimus E*) (*20, 21*) and is also in the Funestus group, somewhat closely related to *An. funestus*, an important African malaria vector. Only *An. minimus s.s*. has been reported from collections in Cambodia (*5*), and is considered to have greater indoor and human biting preferences than other members of this species complex. We currently know very little about the population structure of *An. minimus* and other important vectors occurring across Southeast Asia.

Cambodia is transitioning from focusing on malaria control to malaria pre-elimination, with a goal of eradicating malaria from the country within the next five years. Drug resistant parasites, prevalent *P. vivax* infections, and outdoor malaria transmission are all factors that complicate the goal of malaria elimination in this country. Despite outdoor biting behaviors of many local vector species, Insecticide-treated nets are still the primary vector control method implemented in the GMS, and do have some protective efficacy against *P. falciparum* infection (*22*). Investigations into baseline rates of insecticide resistance have shown that *An. minimus* from different regions have varying levels susceptibility to DDT and pyrethroid insecticides, with some evidence of resistance in populations in Cambodia, Thailand, Laos, India, and Vietnam (*23–25*). Additionally, there is very limited regulation of the use of insecticides for public health and agriculture in Cambodia (*26*).

In the context of the spread of parasites that threaten our best and last-line defence against malaria, we need to know what the population history of the vectors of these parasites look like across this same space. In this study we seek to describe the population structure, diversity, and divergence of populations of *An. minimus* that were active during the rise and spread of drugresistant parasites in Cambodia. We hope to contribute to a better understanding of malaria transmission in Southeast Asia.

## Results

### Population sampling and sequencing

We generated whole genome sequence data from 302 wild-caught individual *An. minimus* female mosquitoes collected from five different field sites in Cambodia using the Illumina HiSeq 2000 platform with 150bp reads with a target coverage of 30X for each. These collections were done in 2010 in Thmar Da, in Eastern Cambodia, a longitudinal monthly collection over 2014 in two sites in each of Pursat, Preah Vihear, and Ratanakiri provinces, and quarterly collections over 2016 in one site each in Pursat and Preah Vihear province, Cambodia.

### Variant discovery

The methods for sequencing and variant calling closely follow those of the Anopheles gambiae 1000 Genomes project phase 2 (Ag1000G) (*27*). Sequence reads were aligned to the *An. minimus* reference genome AminM1 (*28*). The median genome-wide coverage was 35X. We discovered 38,000,285 segregating single nucleotide polymorphisms (SNPs) that passed all of our quality control filters. 13.4% of these SNPs were mutiallelic, with 4,807,355 triallelic and 286,459 quadriallelic SNPs. We restricted our analysis to the largest 40 contigs, which cover 96.6% of the AminM1 reference genome, as many smaller sized contigs can confound diversity and divergence calculations. We found that 75.4% of sites within these largest contigs pass our site filters and thus were accessible to SNP calling.

### Population structure

A principal component analysis (PCA) over biallelic SNPs distributed over the genome of 302 individual field-collected mosquitoes showed that there is clear population structure of *An. minimus* in Cambodia. Samples collected from 5 sites in three provinces split into three distinct populations (Figure 1). One population includes all samples from the western collection site Thmar Da and the northern collection sites in Preah Vihear province, with two further populations with samples from Ratanakiri province in the northeast. These populations split primarily along the first and second principal components. This was a surprising finding because this population structure did not correlate to the geographic sampling of these mosquitoes. Individuals collected from the western and northern sites cluster tighly together despite being hundreds of kilometers apart.

**Figure 1.**
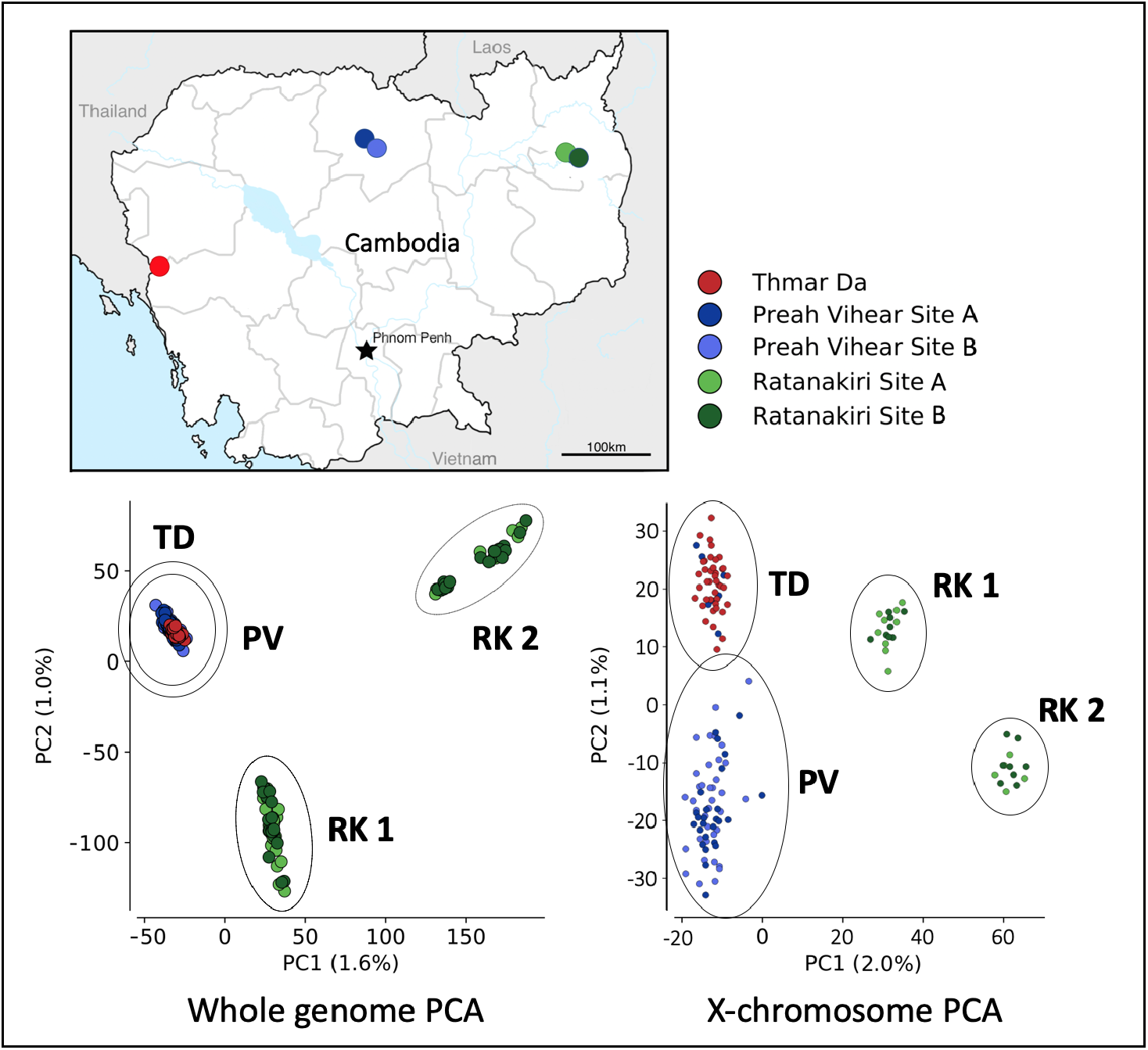
Population structure of *An. minimus* in Cambodia. The map in the top left indicates the five Cambodian collection sites. Principal component analysis (PCA) of whole genome sequences of 302 individual *An. minimus s.s*. collected in five villages in Cambodia shows that there is distinct population structure and three populations. When performing the same PCA on a large X-chromosomal contig (KB664054), these individuals break into four populations: **TD** from the West, **PV** from the northern province in Preah Vihear, and **RK1** and **RK2**, both collected in two sites in Ratanakiri province in the Northeast.

To further explore this structure, we performed the same PCA over contigs from different regions of the genome, PCA over the largest X-chromosomal contig revealed that there is more structure within this quickly evolving sex chromosome, with a splitting of the western and northern samples, indicating four distinct populations of *An. minimus* in Cambodia (Figure 1). The populations defined by these PCA clusters are designated in this study as **TD** from Thmar Da, in Western Cambodia (n=41), on the Thai-Cambodian border, **PV** from the Northern province Preah Vihear (n=156), and the two distinct populations collected in Ratanankiri province in the Northeast, each including individuals collected over multiple years and at both collection sites, these are designated as populations **RK1** (n=58) and **RK2** (n=28).

To examine population differentiation, we computed differences in allele frequencies between each population using Pairwise Fst. Pairwise Fst between all 4 populations over the largest contig, KB663610, representing 16% percent of the *An. minimus* genome, (Figure 2) shows that differentiation was relatively low between populations of TD and PV with an average pairwise Fst of 0.003, while the difference between RK2 and the other three populations is ten-fold higher, around 0.03. Pairwise Fst estimates comparing these populations over other large *An. minimus* contigs indicates a similar level of differentiation, with average pairwise Fst values over 0.03 (Supplementary Table 3). The two sympatric populations from the Ratanakiri collection sites are as differentiated from eachother as they are from the northern and western cluster.

**Figure 2.**
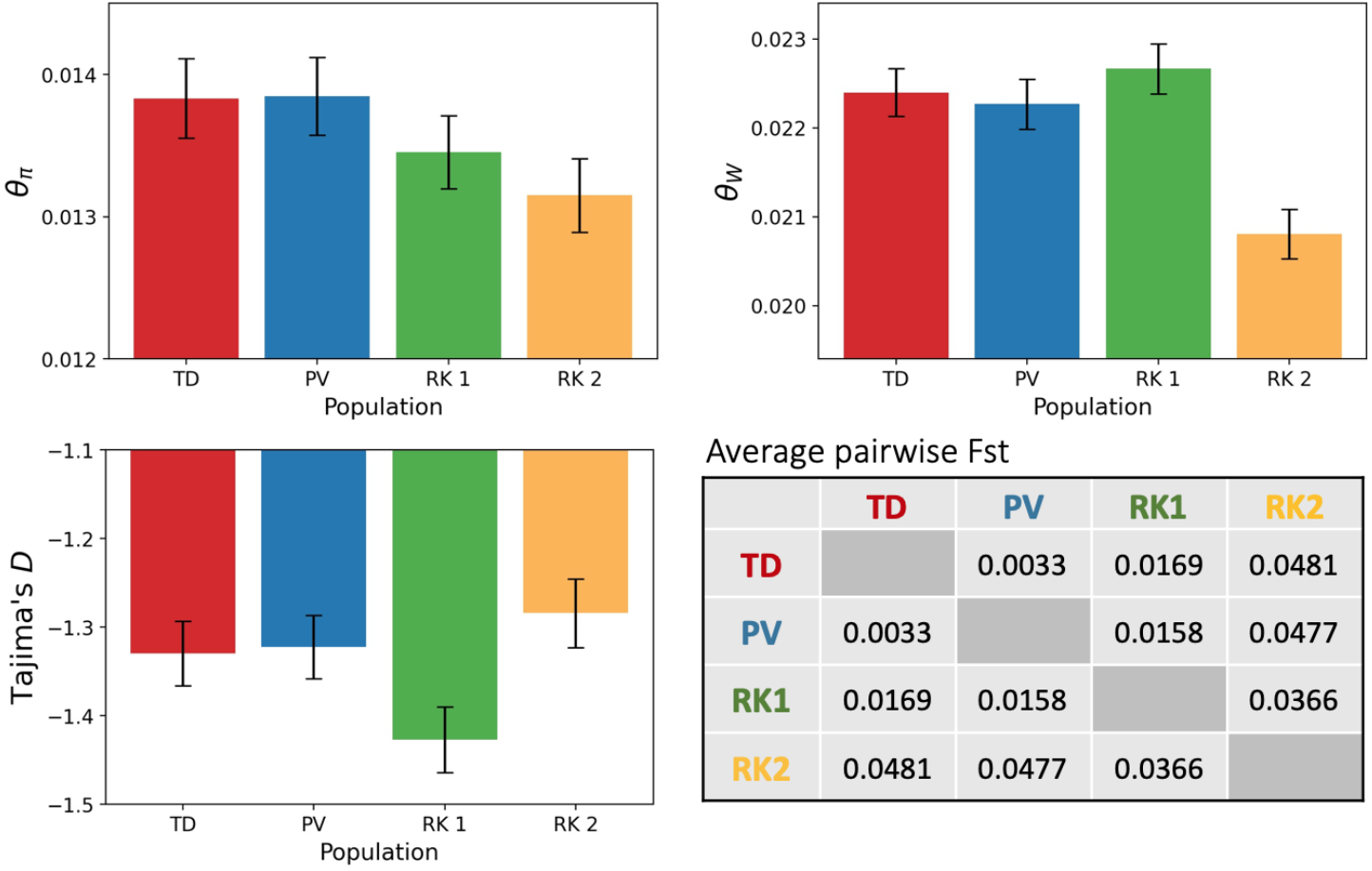
Population diversity and divergence. Nucleotide diversity (π), Watterson’s Theta (θ_W_), and Tajima’s D statistics were calculated over 4-fold degenerate sites on autosomal contigs. The error bars indicate 95% confidence intervals calculated over 100 bootstrap replicates over samples. An average pairwise Fst in the table here was calculated in 20Kb windows over the largest contig KB663610.

This level of differentiation of RK2, even from the RK1 population, might indicate an emerging cryptic species within *An. minimus* A or a newly diverging clade. RK1 and RK2 are sympatric populations, both being collected in the same two sites in Northeastern Cambodia. The differences seen here between RK1 and RK2 populations is consistent with cryptic taxa in other anopheline groups. For example, in the *An. gambiae* complex, the level of differentiation between recently diverged sibling species *An. coluzzii* and *An. gambiae* in West Africa is approximately 0.03 (*19*).

### Population diversity and variation

To characterize population diversity among these populations, nucleotide diversity (π), Watterson’s Theta (θ_W_), and Tajima’s D statistics were calculated over 4-fold degenerate sites on autosomal contigs larger than 2 megabases with 100 bootstrap replicates over samples. These 17 contigs represent 80% of the *Anopheles minimus* genome (Figure 2). The populations were downsampled for these calculations to have same sizes equal to that of the smallest population RK2 (n=28).

There are small but significant differences in the magnitude of the genetic diversity summary statistics between these four different populations. In particular, there were notable differences between the putatively cryptic taxa RK1 and RK2, two populations that were collected in the same sites. RK1 had higher levels of nucleotide diversity and lower levels of Tajima’s D than RK2. These differences are consistent with different population size history between these sympatric groups. In particular, higher levels of genetic diversity indicate larger effective population sizes of TD and PV. Lower values of Tajima’s D suggest stronger population growth in RK1.

These calculations indicate that RK2 has a significantly reduced nucleotide diversity and Watterson’s Theta compared to the other three populations. This may indicate a smaller population size and recent bottleneck of the RK2 population in Cambodia. All four *An. minimus* populations have a negative Tajima’s D, indicating an excess of rare variants, particularly in RK1. This suggests recent population expansions in all populations.

### Signals of evolutionary selection

We used Fst to scan across the *Anopheles minimus* genome to look for regions of the genome with increased differentiation. When we scanned the genome using pairwise Fst, there were no apparent long signals of differentiation that might indicate a large inversion or other structural variant, known to be major drivers of adaptive evolution in other *Anopheles* groups. To further investigate potential large regions of increased differentiation, we performed scans of nucleotide diversity (π), Watterson’s Theta (θ_W_), and Tajima’s D over the largest 14 contigs (representing 80% of the *An. minimus* genome). As with the Fst scans, there were not large regions of higher differentiation in any of the populations that might indicate major structural variants or inversions (Supplementary Figures 1-3).

Whole genome sequencing allowed us to identify pointed signals occurring across the entire genome using scans of average pairwise Fst. Isolated points of high differentiation were compared over single contigs with average pairwise Fst calculated over 1000 SNP windows and plotted over whole contigs. The strongest signals, indicated by the highest Fst value at the peak of a strong signal of differentiation, were ranked and compared. The five top signals in each of six comparisons between the four populations are listed in Table 1. These isolated points of high differentiation are one indication of a signal of evolutionary selection. The most differentiated regions by Fst occurred when comparing the RK2 population to the other three populations, with the highest selection peaks with pairwise Fst over 0.4. RK2 also had more distinct signals of selection when compared to the other populations than RK1. Since these signals of differentiation were highly localized, we could look to known gene annotations and gene predictions across the AminM1 reference genome to see which genes were close to the peaks of these signals. We have noted candidate genes of interest that were very close to the strongest Fst signal peaks and also had known or predicted gene functions (Table 1).

**Table 1.** The top five Fst signals of high differentiation within each of six population comparisons are reported here. Signals were identified using Fst scans in 1000 SNP windows over AminM1 contigs that were larger than 2Mb. Genes and gene predictions near the signal peaks were identified using the VePathDB genome browser.

| Population Comparison | Contig | Signal Position (Mb) | Fst Value | Candidate genes in region | gene(s) | gene functions |
| --- | --- | --- | --- | --- | --- | --- |
| **RK1 v RK2** | KB663622 | 1.25 | 0.54 | UDP-glucuronosyltransferase | AMIN004611 | pyrethroid resistance, detoxification |
| **RK1 v RK2** | KB663622 | 2.2 | 0.52 | Metalloendopeptidase, aminopeptidase N | AMIN004703, AMIN004700, AMIN004701 | pyrethroid resistance, decreased susceptibility to BT toxin |
| **RK1 v RK2** | KB664266 | 6 | 0.49 | Ecdysteroid UDP-glucosyltransferase | AMIN009683 | pyrethroid resistance |
| **RK1 v RK2** | KB663832 | 5.8 | 0.49 | histone acetyltransferase, α-tubulin, CHK domain-containing protein | AMIN016061, AMIN000390, AMIN000391 | dieldrin resistance, organophosphate interactions, mitochondrial function |
| **RK1 v RK2** | KB663610 | 2 | 0.48 | Peptidase S1 domain-containing protein | AMIN002286 | immune response to parasites, detoxification |
| **PV v RK2** | KB663622 | 1.25 | 0.54 | UDP-glucuronosyltransferase | AMIN004611 | pyrethroid resistance, detoxification |
| **PV v RK2** | KB664277 | 0.4 | 0.53 | COP9 signalosome complex subunit | AMIN001615 | hormone and neural signalling |
| **PV v RK2** | KB663610 | 0.1 | 0.53 | cyclic AMP-responsive element-binding protein, JNK1/MAPK8-associated membrane protein | AMIN002152, AMIN002155 | G protein-coupled receptor (GPCR) signal activated transcription factor, stress response |
| **PV v RK2** | KB664266 | 6 | 0.51 | Ecdysteroid UDP-glucosyltransferase | AMIN009683 | pyrethroid resistance |
| **PV v RK2** | KB663622 | 2.2 | 0.5 | Methionine aminopeptidase | AMIN004701 | pyrethroid resistance, decreased susceptibility to BT toxin |
| **PV v RK1** | KB663832 | 1.4 | 0.39 | Elongation of very long chain fatty acids protein | AMIN000203 | lipid concentration, metabolic detoxification |
| **PV v RK1** | KB663832 | 3.7 | 0.38 |  |  | mixed, unspecified products |
| **PV v RK1** | KB664054 | 2.6 | 0.34 | NADPH--cytochrome P450 reductase, 39S ribosomal protein L22 | AMIN005245, AMIN005257 | insecticide resistance, oxidative stress |
| **PV v RK1** | KB663622 | 1.8 | 0.33 | csk homologous kinase domain-containing protein, translation initiation factor 1A | AMIN004659, AMIN004663 | JH signalling, hemocyte activation |
| **PV v RK1** | KB663655 | 1 | 0.33 |  |  | mixed, unspecified products |
| **TD v RK2** | KB663610 | 0.1 | 0.55 | cyclic AMP-responsive element-binding protein, JNK1/MAPK8-associated membrane protein | AMIN002152, AMIN002155 | G protein-coupled receptor (GPCR) signal activated transcription factor, stress response |
| **TD v RK2** | KB663832 | 1.3 | 0.55 | sodium-independent sulfate anion transporter, Protein arginine methyltransferase RmtB, Phosphate carrier- mitochondrial | AMIN000191, AMIN015783, AMIN015787 | signal transduction, phosphate carrier activated in mosquitoes by malaria infection |
| **TD v RK2** | KB663622 | 1.25 | 0.52 | UDP-glucuronosyltransferase | AMIN004611 | pyrethroid resistance, detoxification |
| **TD v RK2** | KB664266 | 6 | 0.5 | Ecdysteroid UDP-glucosyltransferase | AMIN009683 | pyrethroid resistance |
| **TD v RK2** | KB664277 | 0.4 | 0.49 | COP9 signalosome complex subunit | AMIN001615 | hormone and neural signalling |
| **TD v RK1** | KB663832 | 1.25 | 0.5 | UDP-glucuronosyltransferase | AMIN004611 | pyrethroid resistance, detoxification |
| **TD v RK1** | KB664054 | 2.6 | 0.49 | NADPH--cytochrome P450 reductase, 39S ribosomal protein L22 | AMIN005245, AMIN005257 | insecticide resistance, oxidative stress |
| **TD v RK1** | KB663832 | 3.75 | 0.4 |  |  | mixed, unspecified products |
| **TD v RK1** | KB663622 | 1.8 | 0.34 | CHK domain-containing protein, translation initiation factor 1A | AMIN004659, AMIN004663 | mitochondrial function |
| **TD v RK1** | KB663622 | 3.1 | 0.32 | Carbohydrate sulfotransferase | AMIN004763 | detoxification |
| **TD v PV** | KB663622 | 3.1 | 0.125 | Carbohydrate sulfotransferase | AMIN004763 | detoxification |
| **TD v PV** | KB663832 | 0.6 | 0.049 | Carboxylesterase | AMIN000169 | pyrethroid resistance and detoxification |
| **TD v PV** | KB664054 | 2.7 | 0.042 | NADPH--cytochrome P450 reductase, 39S ribosomal protein L22 | AMIN005245, AMIN005257 | insecticide resistance, oxidative stress |
| **TD v PV** | KB663666 | 0.5 | 0.032 | solute carrier family 6 neurotransmitter transporter | AMIN009210 | neural signalling |
| **TD v PV** | KB663832 | 1.35 | 0.027 | sodium-independent sulfate anion transporter, Protein arginine methyltransferase RmtB, Phosphate carrier- mitochondrial | AMIN000191, AMIN015783, AMIN015787 | signal transduction, phosphate carrier activated in mosquitoes by malaria infection |

There is almost no indication of selection when comparing the Thmar Da population with Preah Vihear, with all but one signal with Fst values below 0.05. The one strong signal between TD and PV (Fst = 0.125) is near a Carbohydrate sulfotransferase, which is involved in detoxification processes. Comparing TD to RK1 and RK2 reveals multiple strong signals of selection, some that are present in both Northeastern populations as well as many unique RK2-specific signals (Figure 3, Supplementary figure 4).

**Figure 3.**
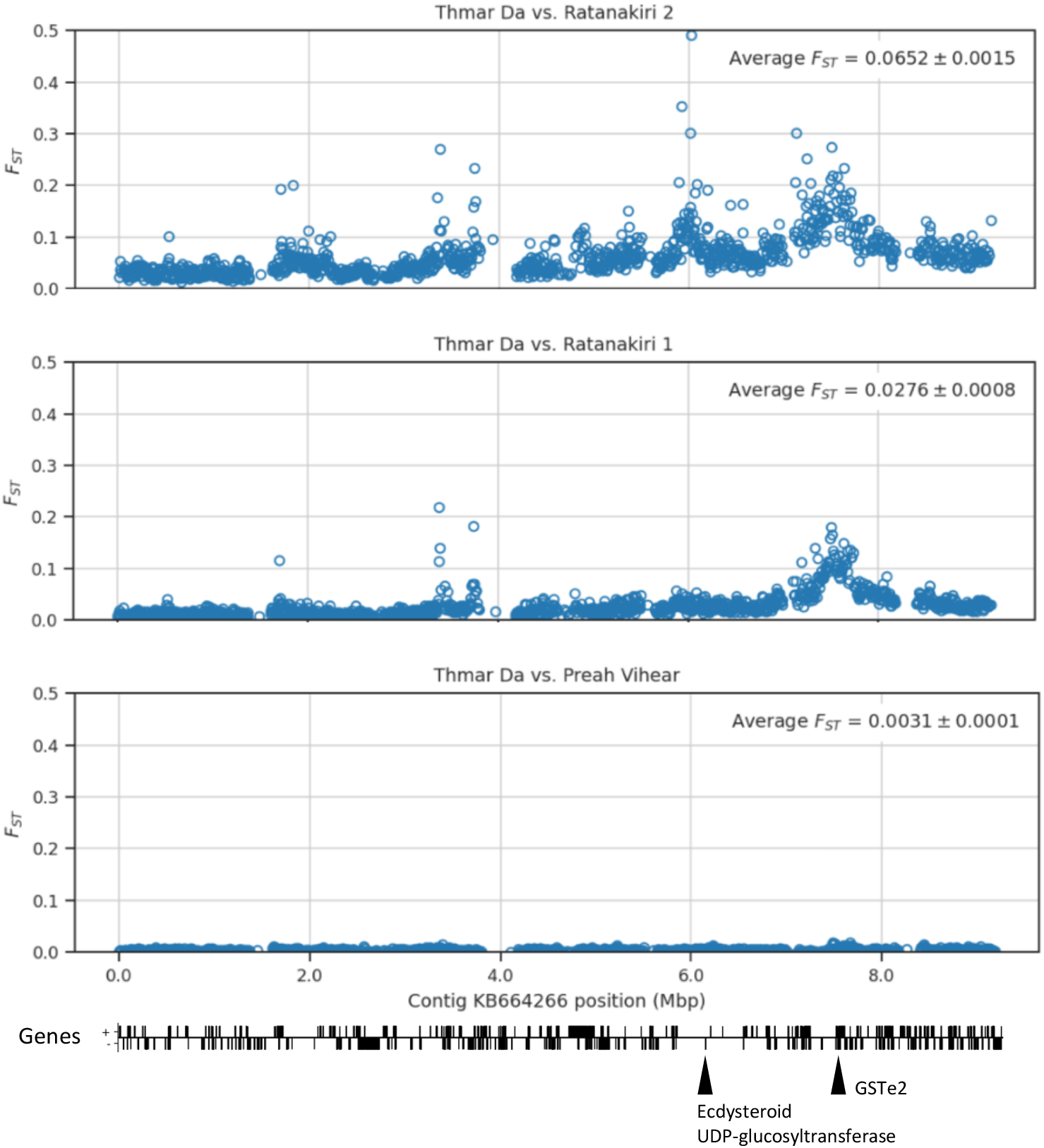
Signals of selection over a single autosomal contig, comparing the Thmar Da population to the three other populations, Ratanakiri 2, Ratanakiri 1, and Preah Vihear. There is almost no indication of selection when comparing Thmar Da with Preah Vihear. There is a strongly supported signal in both Ratanakiri 1 and Ratanakiri 2 populations at 7.5Mb, which is in the same location as a cluster of GSTe genes, including GSTe2, which are known to be involved in metabolic resistance to DDT and pyrethroids. The signal with the highest Fst peak here in RK2, at 6Mb is close to an Ecdysteroid UDP-glucosyltransferase gene, shown to confer pyrethroid insecticide resistance in other anophelines. These are a few of many selection signals identified in this study which may be associated with insecticide pressure on these *An. minimus* populations.

Many of the strongest signals identified in this study may be associated with insecticide pressure on these *An. minimus* populations. The strongest selection signals in every population comparison were close to genes that are involved in detoxification, signal transduction, adaptations to oxidative stress, or have been functionally validated to have mutations that confer resistance to insecticides (Table 1). Some signals of interest include a strongly supported signal of selection in both RK1 and RK2 populations at 7.5Mb on the contig KB664266, which is in the same location as a cluster of glutathione-S-transferases, including GSTe2, which has been shown to be involved in the metabolism of DDT and pyrethroids, mutations in which mediate metabolic insecticide resistance (*29*). The signal with the highest pairwise Fst peak on the same contig KB664266, at 6Mb is an RK2 specific signal and close to an Ecdysteroid UDP-glucosyltransferase gene, which has been shown to confer pyrethroid insecticide resistance in *An. stephensi* (*30*).

Another notable signal is between the RK1 and RK2 populations on the contig KB663610, a Peptidase S1 domain-containing protein AMIN002286, which has been shown to be involved in response to parasite pathogens in insects (*31*). The signals of selection observed in this study are mostly distinct from the main selection signals seen in *An. gambiae* complex mosquitoes (*19*), the primary vectors of *Plasmodium falciparum* in Africa.

#### Insecticide resistance

We report here variants at known insecticide resistance associated alleles for each of the four *An. minimus* populations. Variants occurring at a frequency of more than 2% in at least one of the four populations are reported in the known insecticide-resistance associated genes Ace1, Rdl, KDR, and GSTe2 (Supplementary Table 2). GSTe2 mutants are present in multiple populations, at a low rate, and there are a few individuals in Thmar Da and Preah Vihear with the Rdl resistance mutation, which is known to confer resistance to cyclodiene insecticides, despite evidence from other studies that species in this region lack this resistance mutation (*32*). We did not investigate copy number variation, which is one mechanism by which GSTe2 confers insecticide resistance. These SNP variants indicate variation throughout these insecticide-resistance associated genes and though most of these population do not currently have high rates of validated insecticide resistance associated mutations, this underlying variation provides potential for structural and transcriptional events resulting in greater levels of insecticide resistance in *An. minimus* populations.

## Discussion/Conclusions

This is the first study investigating the population genomics of *An. minimus*. We examined the whole genomes of 302 wild-collected female mosquitoes from five different sites in Cambodia over several years. This important vector species appears to be highly structured within Cambodia, with a division of West, North, and North-eastern populations. This structure cannot be fully accounted for by geography. The RK1 and RK2 populations that are as differentiated from each other as *An. coluzzii* is from *An. gambiae s.s.*, were collected in the same sites and at the same timepoints. The other populations PV and TD are separated by a few hundred kilometres with no major geographic barrier between these regions of Cambodia. The sympatric RK1 and RK2 populations are as distinct from one another as they are from the geographically distinct populations PV and TD. The RK2 population likely represents an emerging cryptic species or diverging clade within *An. minimus s.s*.

All four populations appear to be under selective pressure and may have undergone recent population expansions. These diverging populations of *An. minimus* each have unique signals of evolutionary selection. These signals are particularly prominent in the most differentiated population, RK2. Many of the signals identified in this study indicate a strong evolutionary response to insecticidal pressure in the GMS, this pressure is more likely due to heavy and longterm agricultural pesticide use rather than malaria vector control such as indoor residual spray or long-lasting insecticidal nets. Multiple mechanisms of insecticide resistance have been identified in other insects and in other disease vector mosquitoes in Cambodia (*23, 24, 33*). The widespread use of pyrethroid, organophosphate, and carbamate insecticides in agriculture exacerbates the development of insecticide-resistance in populations of many different species across the GMS (*23*).

This study shows that populations of *An. minimus* in Cambodia are evolving, and this may be in the context of strong and sustained pressure from agricultural insecticide use. Each population in this study has unique signals of evolutionary selection associated with multiple detoxification pathways and may also be under different types of insecticidal pressure than mosquitoes in other parts of the world. For example, since *An. minimus* is a species that is able to exploit rice paddies and flooded agricultural fields as larval habitats, individual mosquitoes may experience insecticidal pressure as larvae, which might impact their susceptibility to insecticides as adults. The presence of these signals reveals a complicated landscape of potentially multiple insecticide-resistant vectors that may not be responsive to insecticides implemented as part of a malaria control campaign.

Current vector control tools in Cambodia and most of the GMS include the widespread distribution of long-lasting insecticidal nets (LLINs), and topical repellents. New control tools are needed in this region and in the context of outdoor, residual malaria transmission. The population structure within this single Southeast Asian vector species in a small country like Cambodia may indicate barriers for implementing vector control methods centered around genetic modification or sterile insect release. If populations of vectors are diverging with limited gene flow, despite being in the same place, genetic control based on any assumption of a single population of vectors would be unlikely to penetrate vector populations in this region.

This study expands the *Anopheles* species represented in whole genome population data. This resource of *An. minimus* whole genome data will be a starting point to explore *Anopheles* genome variation in important transmission contexts such as Southeast Asia. This study is the foundation for further delving into the population structure and history of *An. minimus*, which acts as an important malaria vector across all of Southeast Asia.

Examining the population structure in a single species over its range and through time will help us to better understand the potential risk associated with multiple insecticide-resistant populations of vectors and how to best approach managing and monitoring insecticide resistance. *An. minimus* in Cambodia includes an adaptive, resilient, and diverse set of populations that can thrive in human-created habitats and potentially contribute to epidemics of drug-resistant malaria as we work toward malaria elimination. It is crucial that we appreciate the evolutionary history and ecology of this important and widespread malaria vector.

## Materials and Methods

### Collection methods

*An. minimus* female mosquitoes were selected from several previous field collections in Cambodia with the goal of including a subsample of *An. minimus* specimens from different field sites and timepoints for population genomic analysis. These anophelines were collected using CDC light traps, human landing collections, cow-baited tents, and barrier fences from seven different locations in Cambodia, from 2009 to 2014. Five of these sites had enough *An. minimus* individuals to include in population genomic analyses.

Collection locations included Thmar Da, in Western Cambodia on the Thai/Cambodian border (Thmar Da) and in Pursat Province (Angkrong and Veal villages), in the North in Preah Vihear province (Chean Mok pagoda and Preah Kleang village), and in the Northeast Ratanakiri (Sayas and Chamkar Mai villages)(*34, 35*). Multiple *Anopheles* species were collected in each of these studies, including the *An. minimus A* specimens that have been included in this study. GPS coordinates for each collection site are available in the sample metadata (Supplementary Table 1, 5).

### DNA extraction and identification

Specimens were morphologically identified immediately after collection using keys from (*36, 37*). These were stored individually in 1.5ml tubes with silica gel desiccant. DNA was extracted from the head and thorax of collected mosquitoes using either a CTAB extraction method or in Nextec plates. Individual mosquitoes were molecularly identified using Sanger sequencing of the rDNA ITS2 region using primers ITS2A and ITS2B from (*38*) and compared to voucher reference rDNA ITS2 sequences in NCBI.

### Whole genome Sequencing

Individual samples were sequenced at the Wellcome Trust Sanger Institute using whole-genome sequencing on the Illumina HiSeq 2000 platform. Paired-end multiplex libraries were prepared as per the manufacturer’s specifications, with DNA fragmented using Covaris Adaptive Focused Acoustics. Multiplexes comprised 12 tagged individual mosquitoes. Three lanes of sequencing were used for each multiplex to control for variation between sequencing runs. Sequencing and cluster generation were done using the manufacturer’s protocol for paired-end reads of 150bps with insert sizes from 100-200 bp. The target coverage for each individual was 30X. To include some samples that had DNA concentration that was too low to run through the standard Illumina pipeline, we whole genome sequenced a set of specimens using a new low-input DNA sequencing pipeline (42). This low-input sequencing pipeline uses enzymatic fragmentation and less than 10 nanograms of input DNA. In our sequence and population QC, these low-input samples did not appear to be of lower quality or influence the clustering of whole genome sequences in our PCA analysis. 144 individual *An. minimus* mosquitoes were sequenced using the standard Illumina pipeline and 158 individuals were sequenced using the low-input pipeline.

### Sequence alignment

Illumina whole genome sequence reads were aligned to the *Anopheles* reference genome AminM1 using BWA version 0.7.15.(*39, 40*) using bwa mem. A BAM file was constructed for each individual using merged alignments from multiple lanes. Duplicate reads were marked using Picard version 2.6.0 (*41*).

### Coverage

For each sample, depth of coverage was computed at all genome positions. Samples were excluded if median coverage across all chromosomes was less than 10×, or if less than 50% of the reference genome was covered by at least 1×.

### Population outliers and anomaly detection

We used principal component analysis (PCA) to identify and exclude individual samples that were population outliers. SNPs were down-sampled to use 100,000 segregating non-singleton sites from autosomal contigs. PCA was computed using scikit-allel version 1.2.0. We iteratively identified and excluded any individual samples that were outliers along a single principal component. We then identified and excluded any individual samples or small sample groups that clustered together with other samples in a way that was not plausible given metadata regarding their collection location.

### Quantification of population diversity and divergence

Diversity statistics Pairwise Fst, nucleotide diversity, Tajima’s D, and Watterson’s Theta were calculated in windows across the largest 40 contigs on the *An. minimus* AminM1 genome using scikit-allel version 1.3.3. and custom code in Python.

Nucleotide diversity (π), Watterson’s Theta (θ_W_), and Tajima’s D statistics were calculated over 4-fold degenerate sites on autosomal contigs larger than 2,000,000 bases. These 17 contigs represent 80% of the *Anopheles minimus* genome. These statistics were calculated as an average over sliding windows with 100 bootstrap replicates over samples. Populations were downsampled to the size of the smallest population RK2 (n=28) for these calculations.

### Quantification of signals of selection

We used pairwise Fst in 1000 SNP windows to scan across the *Anopheles minimus* genome to look for any signals of increased differentiation, which might indicate evolutionary selection. This scan was performed for contigs larger than 2 Megabases, these contigs represent 86% of the *An. minimus* genome. Signals that were represented by at least 5 1000 SNP windows above two times the genome-wide average were recorded and ranked by the highest Fst value at the peak of each selection signal.

Genes potentially associated with these signals were identified by going to the location of the peak of each signal on the *An. minimus* genome (AminM1) using the Genome Browser tool in VEuPathDB Vectorbase (*28*) and walking in both directions from the location signal to see which annotated genes were present in that location. Gene IDs and gene functional predictions were recorded.

### Variants in known insecticide-resistance associated genes

We report in this study variants at known insecticide resistance associated alleles. Variants occurring at a frequency of more than 2% in at least one of the four populations are reported in the known insecticide-resistance associated genes Ace1, Rdl, KDR, and Gste2 (Supplementary Table 2).

## Supporting information

Supplemental materials summary and figs

Supplementary figures population comparisons

## Data accessibility

These genome and SNP data are available at https://www.malariagen.net/resource/35. These data are hosted in Google Cloud. For more information about downloading data, please see the data download guide at https://malariagen.github.io/vector-data/amin1/intro.html. For more information about accessing data in the cloud, please see the cloud access guide at https://malariagen.github.io/vector-data/amin1/download.html.

## Author contributions

BSTL, AM, and DK conceived of the study. BSTL designed and directed the field collections. BSTL, KO, CS, MS, RS, and SN carried out the field collections in multiple sites in Cambodia with KO managing entomological data collection and SS managing personnel and site coordination. BSTL and ND performed molecular analysis for molecular species identification and plasmodium detection. ND, ED, KR, and SG were involved in preparing samples and sample data for whole genome sequencing. BSTL, NH, and AM carried out the population genomic analyses. BSTL prepared the manuscript. All authors have approved the manuscript.

## Acknowledgements

Thank you to our collaborators at the National Center for Parasitology, Entomology, and Malaria Control (CNM) in Phnom Penh, Cambodia, the National Institutes of Health NIAID LMVR, and the MalariaGEN team for their support of this work.

## Works Cited

1. W. Van Bortel et al., Malaria transmission and vector behaviour in a forested malaria focus in central Vietnam and the implications for vector control. Malar J 9, 373 (2010).

2. L. Durnez et al., Outdoor malaria transmission in forested villages of Cambodia. Malar J 12, 329 (2013).

3. F. M. Hawkes et al., Vector compositions change across forested to deforested ecotones in emerging areas of zoonotic malaria transmission in Malaysia. Scientific reports 9, 13312 (2019).

4. H. D. Trung et al., Behavioural heterogeneity of *Anopheles* species in ecologically different localities in Southeast Asia: a challenge for vector control. Tropical medicine & international health: TM & IH 10, 251–262 (2005).

5. W. Van Bortel et al., Eco-ethological heterogeneity of the members of the *Anopheles minimus* complex (Diptera: Culicidae) in Southeast Asia and its consequences for vector control. Journal of medical entomology 41, 366–374 (2004).

6. V. Dev, S. Manguin, Biology, distribution and control of *Anopheles* (Cellia) *minimus* in the context of malaria transmission in northeastern India. Parasites & vectors 9, 585 (2016).

7. A. Vantaux et al., Anopheles ecology, genetics and malaria transmission in northern Cambodia. Scientific reports 11, 6458 (2021).

8. R. E. Harbach, J. B. Gingrich, L. W. Pang, Some entomological observations on malaria transmission in a remote village in northwestern Thailand. Journal of the American Mosquito Control Association 3, 296–301 (1987).

9. P. Somboon, W. Suwonkerd, J. D. Lines, Susceptibility of Thai zoophilic Anophelines and suspected malaria vectors to local strains of human malaria parasites. Southeast Asian J Trop Med Public Health 25, 766–770 (1994).

10. C. Amaratunga et al., Dihydroartemisinin & piperaquine resistance in Plasmodium falciparum malaria in Cambodia: a multisite prospective cohort study. The Lancet Infectious Diseases, (2016).

11. O. Miotto et al., Multiple populations of artemisinin-resistant *Plasmodium falciparum* in Cambodia. Nature genetics 45, 648–655 (2013).

12. S. Takala-Harrison et al., Independent Emergence of Artemisinin Resistance Mutations Among *Plasmodium falciparum* in Southeast Asia. The Journal of infectious diseases 211, 670–679 (2015).

13. W. L. Hamilton et al., Evolution and expansion of multidrug-resistant malaria in southeast Asia: a genomic epidemiology study. The Lancet. Infectious diseases 19, 943–951 (2019).

14. B. St Laurent et al., Artemisinin-resistant *Plasmodium falciparum* clinical isolates can infect diverse mosquito vectors of Southeast Asia and Africa. Nat Commun 6, 8614 (2015).

15. P. Poolphol et al., Natural *Plasmodium vivax* infections in *Anopheles* mosquitoes in a malaria endemic area of northeastern Thailand. Parasitol Res 116, 3349–3359 (2017).

16. L. Cui et al., Malaria in the Greater Mekong Subregion: heterogeneity and complexity. Acta Trop 121, 227–239 (2012).

17. M. E. Sinka et al., The dominant *Anopheles* vectors of human malaria in the Asia-Pacific region: occurrence data, distribution maps and bionomic precis. Parasites & vectors 4, 89 (2011).

18. M. C. Fontaine et al., Mosquito genomics. Extensive introgression in a malaria vector species complex revealed by phylogenomics. Science 347, 1258524 (2015).

19. C. Anopheles gambiae Genomes et al., Genetic diversity of the African malaria vector *Anopheles gambiae*. Nature 552, 96–100 (2017).

20. WHO, Anopheline Species Complexes in South and South-East Asia. S. K. Subbarao, Ed., (World Health Organization 2007), pp. 102.

21. C. Garros, R. E. Harbach, S. Manguin, Systematics and biogeographical implications of the phylogenetic relationships between members of the funestus and minimus groups of *Anopheles* (Diptera: Culicidae). Journal of medical entomology 42, 7–18 (2005).

22. T. Sochantha et al., Insecticide-treated bednets for the prevention of *Plasmodium falciparum* malaria in Cambodia: a cluster-randomized trial. Tropical medicine & international health 11, 1166–1177 (2006).

23. W. Van Bortel et al., The insecticide resistance status of malaria vectors in the Mekong region. Malaria journal 7, 102 (2008).

24. S. Marcombe et al., Insecticide resistance status of malaria vectors in Lao PDR. PloS one 12, e0175984 (2017).

25. P. Somboon, L. A. Prapanthadara, W. Suwonkerd, Insecticide susceptibility tests of *Anopheles minimus s.l., Aedes aegypti, Aedes albopictus*, and *Culex quinquefasciatus* in northern Thailand. The Southeast Asian journal of tropical medicine and public health 34, 87–93 (2003).

26. H. van den Berg, R. Velayudhan, R. S. Yadav, Management of insecticides for use in disease vector control: Lessons from six countries in Asia and the Middle East. PLoS neglected tropical diseases 15, e0009358 (2021).

27. C. Anopheles gambiae Genomes, Genome variation and population structure among 1142 mosquitoes of the African malaria vector species *Anopheles gambiae* and *Anopheles coluzzii*. Genome Res 30, 1533–1546 (2020).

28. G. I. Giraldo-Calderon et al., VectorBase: an updated bioinformatics resource for invertebrate vectors and other organisms related with human diseases. Nucleic acids research 43, D707–713 (2015).

29. H. Ranson et al., Identification of a novel class of insect glutathione S-transferases involved in resistance to DDT in the malaria vector *Anopheles gambiae*. Biochem J 359, 295–304 (2001).

30. Y. Zhou et al., UDP-glycosyltransferase genes and their association and mutations associated with pyrethroid resistance in *Anopheles sinensis* (Diptera: Culicidae). Malaria journal 18, 62 (2019).

31. X. Cao, M. Gulati, H. Jiang, Serine protease-related proteins in the malaria mosquito, Anopheles gambiae. Insect biochemistry and molecular biology 88, 48–62 (2017).

32. S. Marcombe et al., Malaria and Dengue Mosquito Vectors from Lao PDR Show a Lack of the rdl Mutant Allele Responsible for Cyclodiene Insecticide Resistance. Journal of medical entomology 57, 815–823 (2020).

33. S. Boyer et al., Resistance of *Aedes Aegypti* populations to deltamethrin, permethrinm and temephos in Cambodiain *American Journal of Tropical Medicine and Hygiene*. (AMER SOC TROP MED & HYGIENE 8000 WESTPARK DR, STE 130, MCLEAN, VA 22101 USA, 2018), vol. 99, pp. 265–265.

34. B. St Laurent et al., Cow-baited tents are highly effective in sampling diverse *Anopheles* malaria vectors in Cambodia. Malaria journal 15, 440 (2016).

35. B. St Laurent, Clinically informed mosquito sampling in Cambodia – a year-long survey of diverse and abundant malaria vectors in three provinces. Unpublished.

36. E. Peyton, J. E. Scanlon, V. Malikul, S. Imvitaya, “Illustrated key to the female *Anopheles* mosquitoes of Thailiand,” (Army Medical Component AFRIMS APO San Francisco 96346, 1966).

37. R. Rattanarithikul, B. A. Harrison, P. Panthusiri, E. L. Peyton, R. E. Coleman, Illustrated keys to the mosquitoes of Thailand III. Genera Aedeomyia, Ficalbia, Mimomyia, Hodgesia, Coquillettidia, Mansonia, and Uranotaenia. The Southeast Asian journal of tropical medicine and public health 37 Suppl 1, 1–85 (2006).

38. N. W. Beebe, A. Saul, Discrimination of all members of the *Anopheles punctulatus* complex by polymerase chain reaction--restriction fragment length polymorphism analysis. Am J Trop Med Hyg 53, 478–481 (1995).

39. H. Li, R. Durbin, Fast and accurate short read alignment with Burrows-Wheeler transform. Bioinformatics 25, 1754–1760 (2009).

40. H. Li, R. Durbin, Fast and accurate long-read alignment with Burrows-Wheeler transform. Bioinformatics 26, 589–595 (2010).

41. Broad_Institute, Picard Toolkit. GitHub Repository. http://broadinstitute.github.io/picard/, (2019).

42. Ellis, P., Moore, L., Sanders, M.A. et al. Reliable detection of somatic mutations in solid tissues by laser-capture microdissection and low-input DNA sequencing. Nat Protoc 16, 841–871 (2021).

