## Supplemental materials summary and figs for "Population genomics reveal distinct and diverging populations of *An. minimus* in Cambodia – a widespread malaria vector in Southeast Asia"

**Supplementary Figures:**

Supplementary Figure 1 - Nucleotide diversity across the largest 14 AminM1 contigs for four Cambodian populations

Supplementary Figure 2 - Watterson's Theta across the largest 14 AminM1 contigs for four Cambodian populations

Supplementary Figure 3 - Tajima's D across the largest 14 AminM1 contigs for four Cambodian populations

Supplementary Figure 4 (A-F) - *An. minimus* Fst scan population comparisons
These tables include Fst scans in 1000 SNP windows across the largest 18 AminM1 contigs for each of 6 population comparisons.

**Supplementary Tables** (supplementary_tables_min_pop_gen):

Supplementary Table 1 - sample metadata
This table includes unique sample identifiers and collection metadata for each individual female mosquito included in this study.

Supplementary Table 2 -  IR SNP variants
SNP variants occurring in over 2% in any within known insecticide-resistance associated genes Ace1, Rdl, KDR, and GSTe2, population are reported here.

Supplementary Table 3 - population Fst
Pariwise average Fst calculations in 20Kb windows for the four populations over the five largest contigs are reported.

Supplementary Table 4 – Min contig locations
Summary of largest 40 contigs used for diversity statistics and other calculations in this study, including which *Anopheles* genomic element and AgamP4 chromosome equivalent arm they lie on.

Supplementary Table 5 – sampling summary
A summary of timepoints and sites where individual *An. minimus* samples were collected.

**Supplementary Figure 1** - Nucleotide diversity calculated in 50kb windows across the largest 14 AminM1 contigs for four Cambodian populations

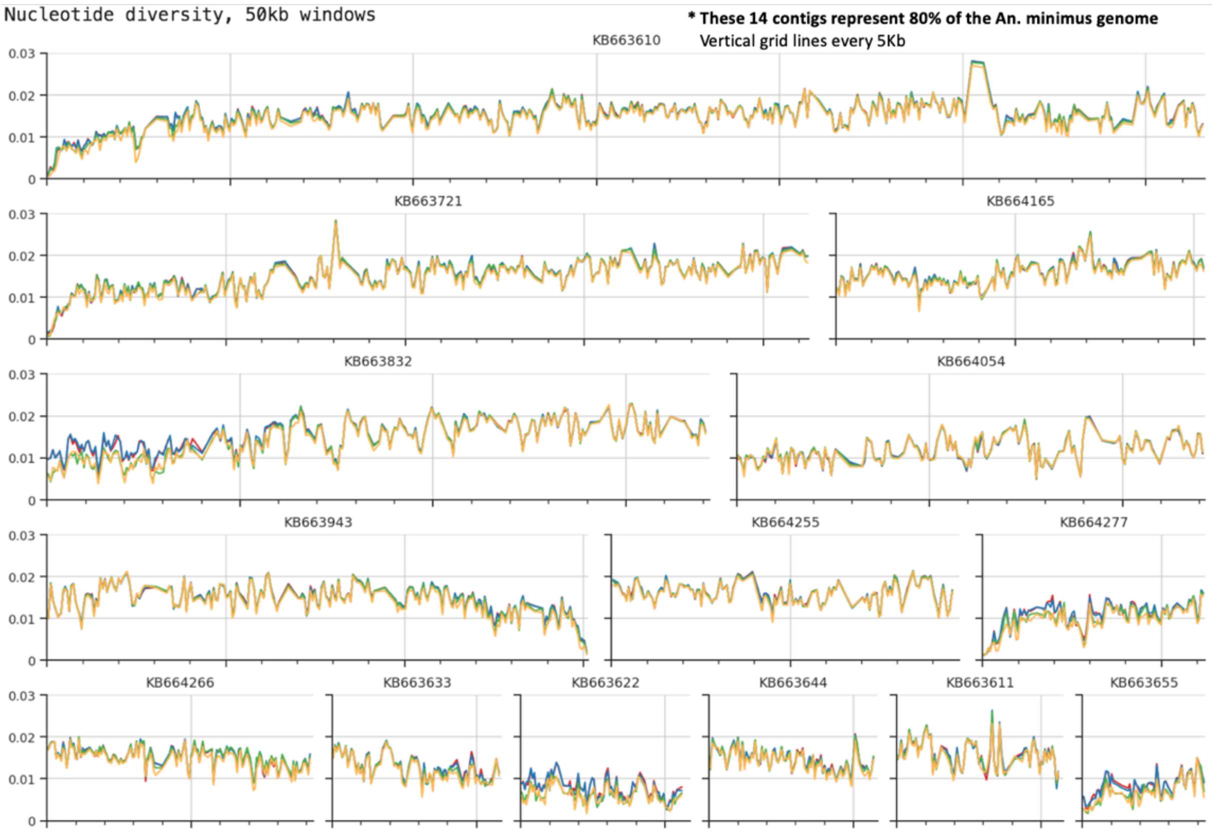

**Supplementary Figure 2** - Watterson's Theta calculated in 50kb windows across the largest 14 AminM1 contigs for four Cambodian populations

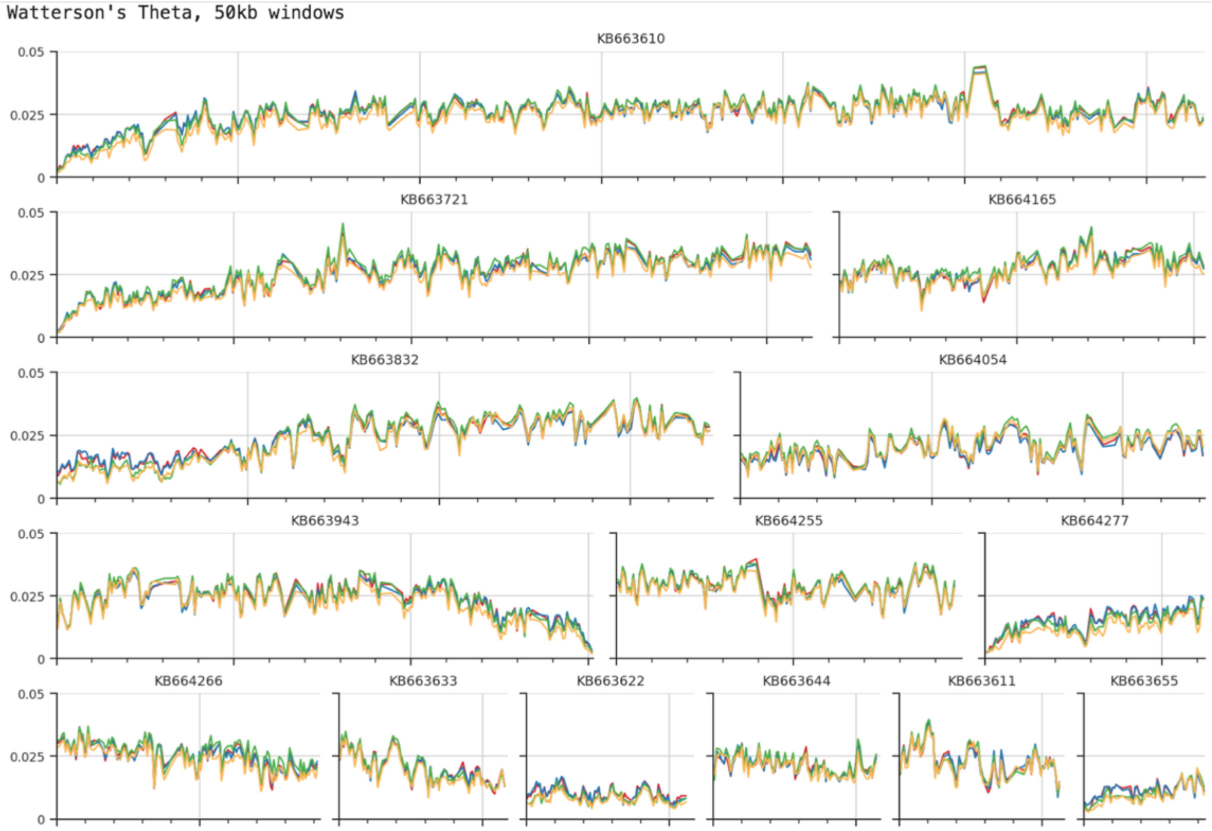

**Supplementary Figure 3** - Tajima's D calculated in 50kb windows across the largest 14 AminM1 contigs for four Cambodian populations

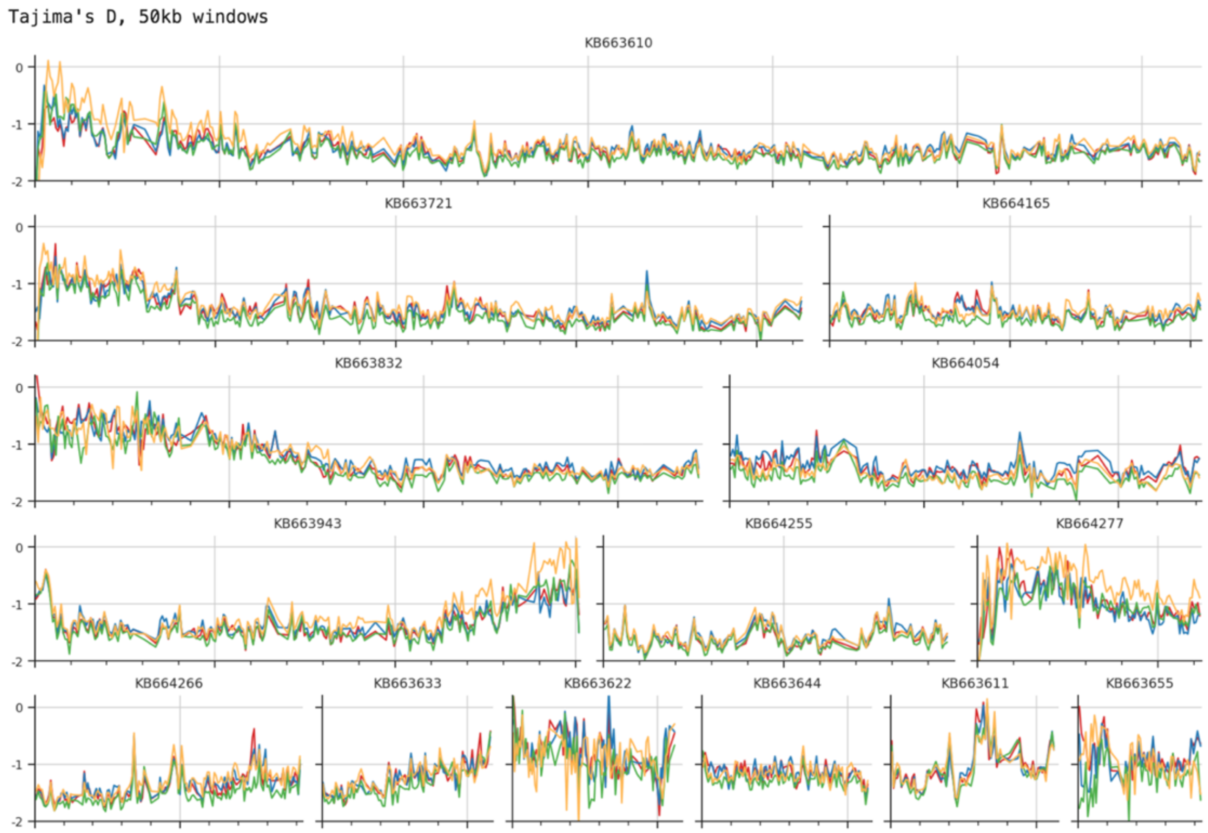

Supplementary Figure 4.1

**TD vs. PV**

Fst scans in 1000 SNP windows across the largest 18 AminM1 contigs

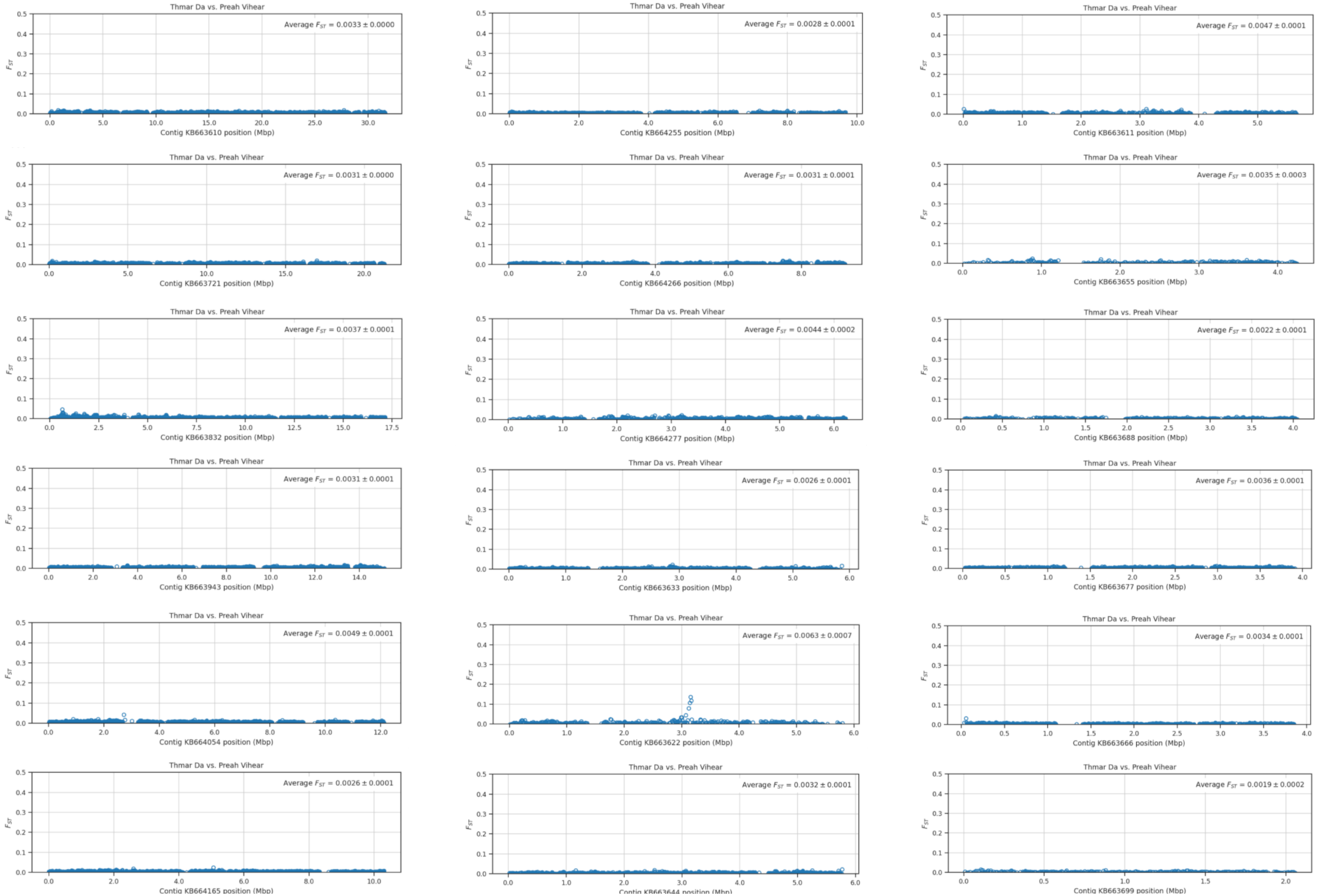

Supplementary Figure 4.2

# TD vs. RK1

Fst scans in 1000 SNP windows across the largest 18 AminM1 contigs

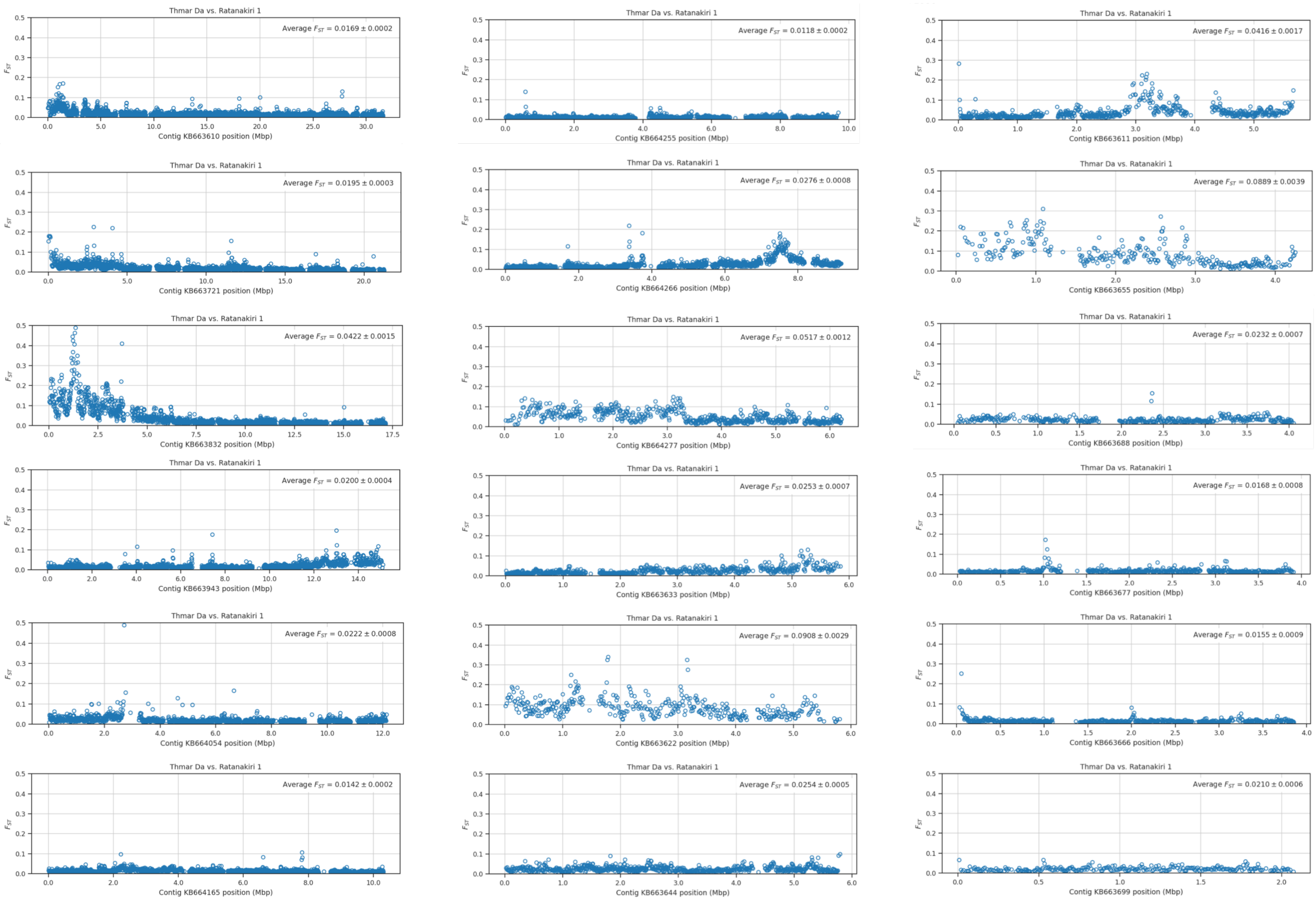

Supplementary Figure 4.3

# TD vs. RK2

Fst scans in 1000 SNP windows across the largest 18 AminM1 contigs

*axes adjusted to accommodate Fst greater than 0.5

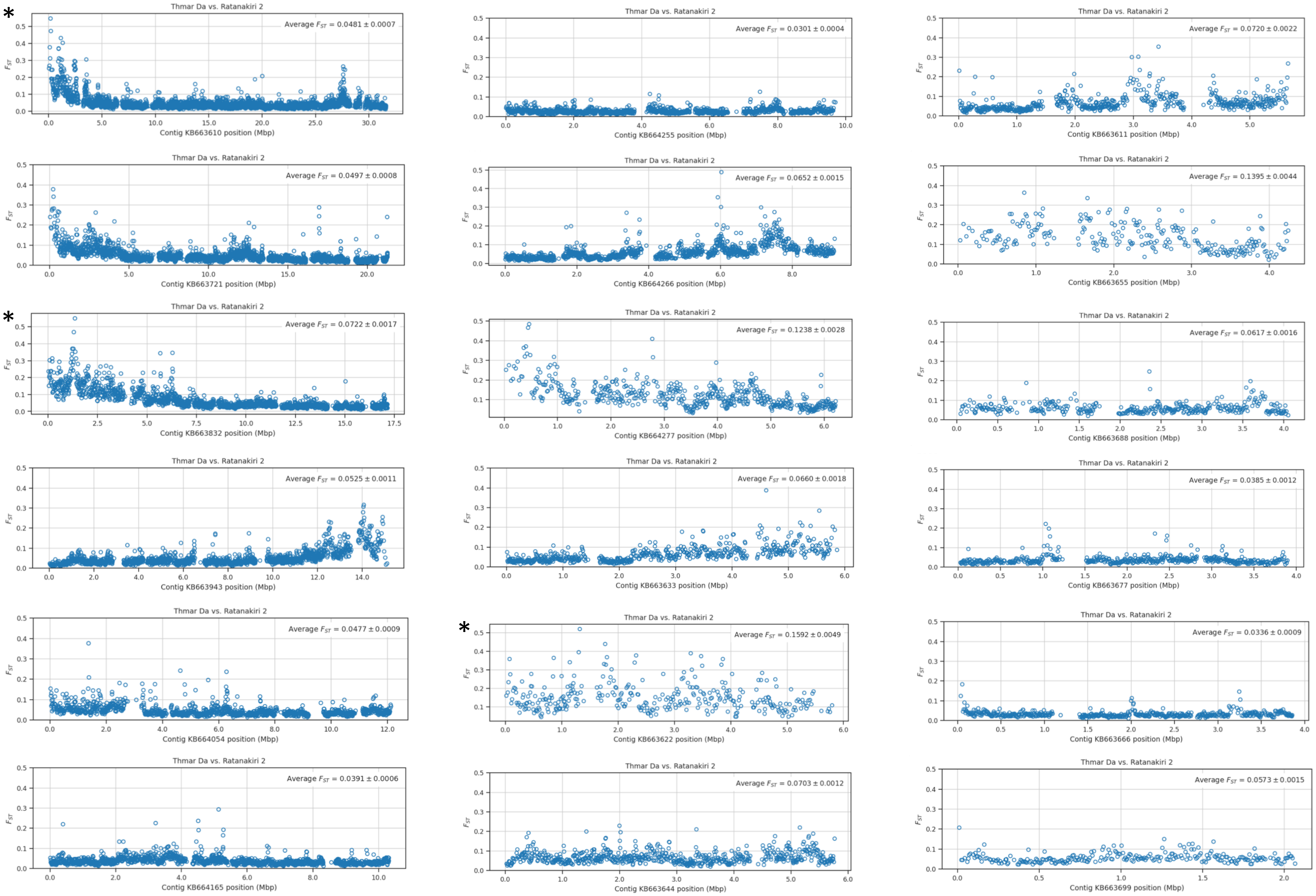

Supplementary Figure 4.4

**PV vs. RK1**

Fst scans in 1000 SNP windows across the largest 18 AminM1 contigs

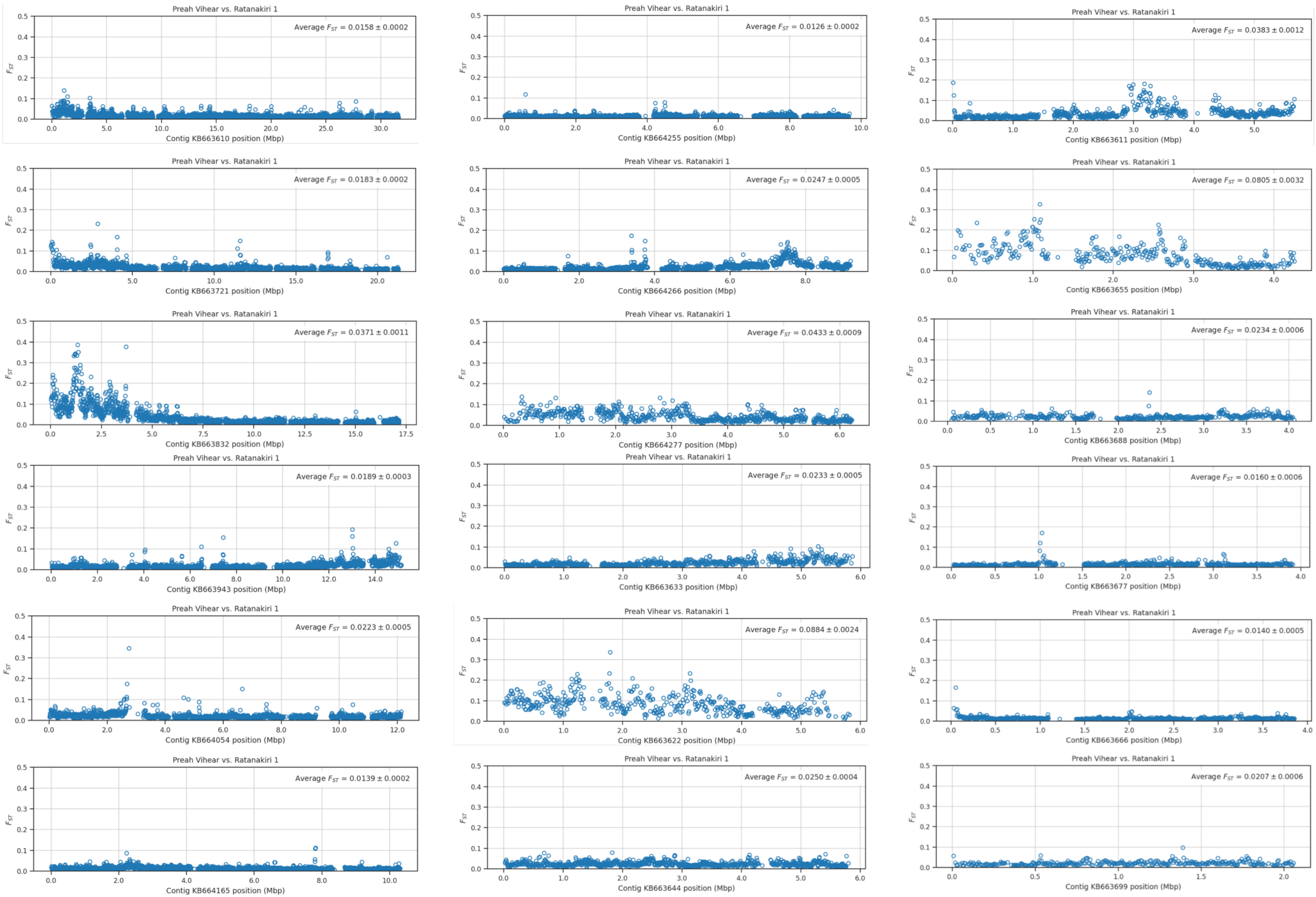

Supplementary Figure 4.5

# PV vs. RK2

Fst scans in 1000 SNP windows across the largest 18 AminM1 contigs

*axes adjusted to accommodate Fst greater than 0.5

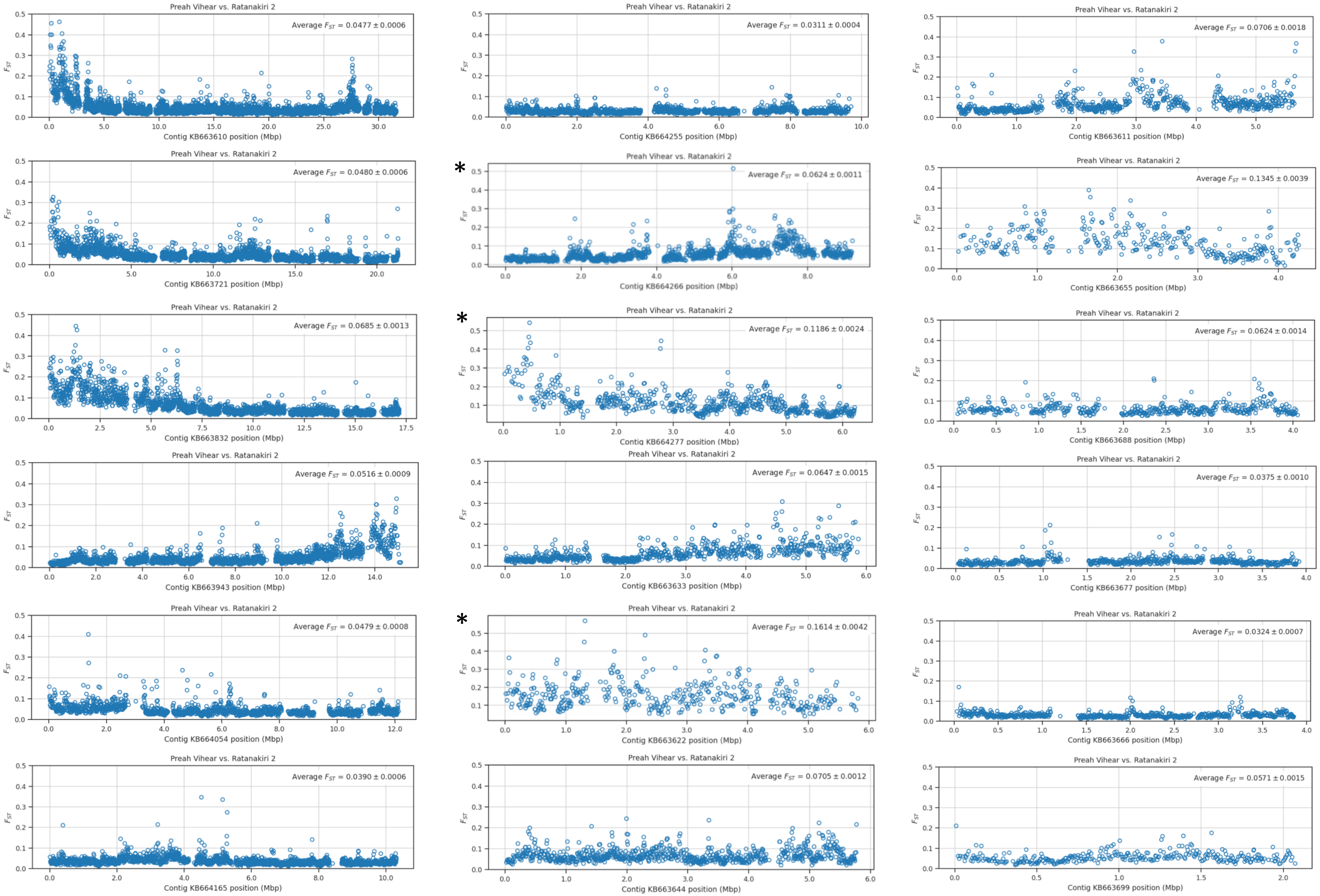

Supplementary Figure 4.6

# RK1 vs. RK2

Fst scans in 1000 SNP windows across the largest 18 AminM1 contigs

*axes adjusted to accommodate Fst greater than 0.5

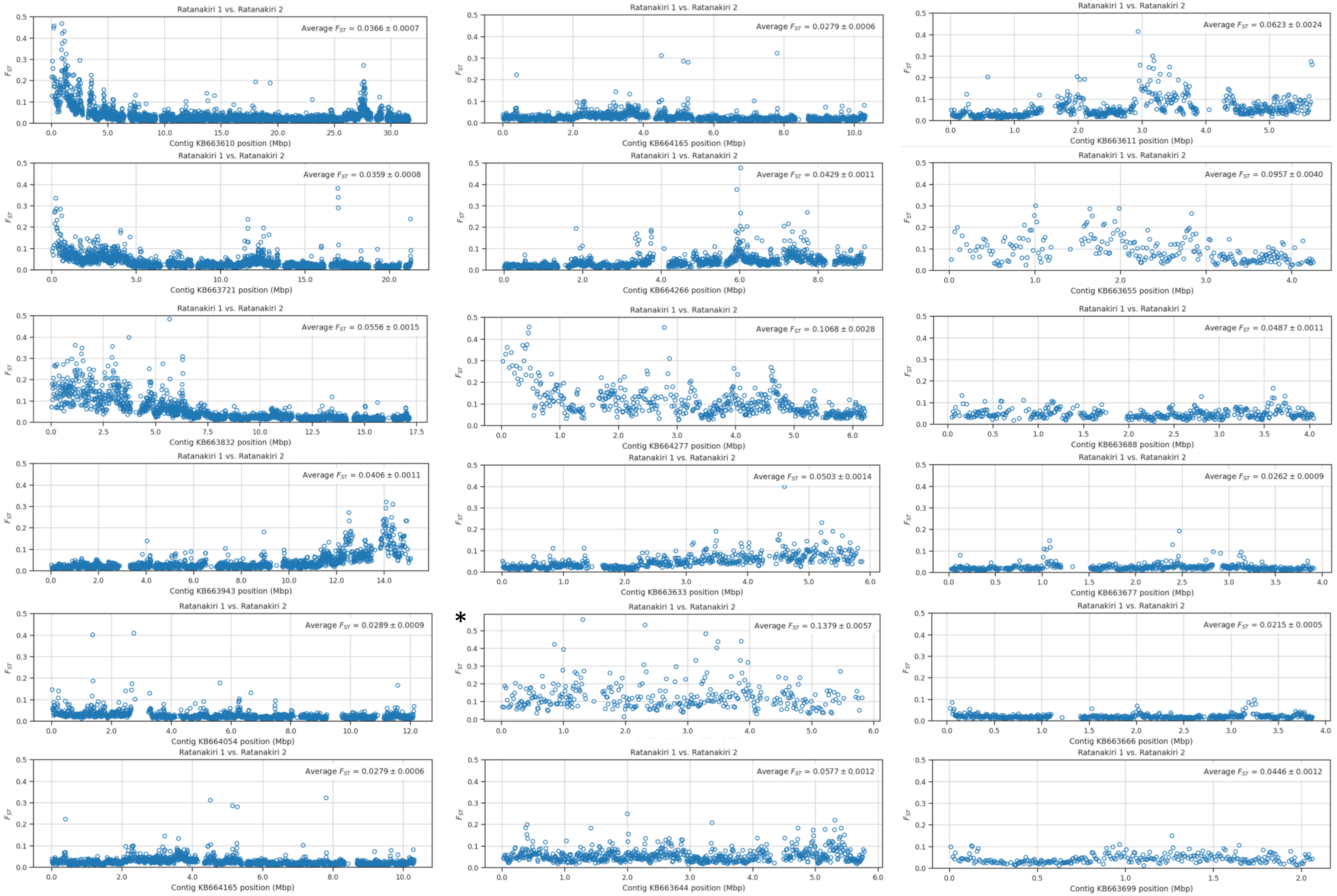
