## Supplementary figures population comparisons for "Population genomics reveal distinct and diverging populations of *An. minimus* in Cambodia – a widespread malaria vector in Southeast Asia"

Supplementary Figure 4.1

**TD vs. PV**

Fst scans in 1000 SNP windows across the largest 18 AminM1 contigs

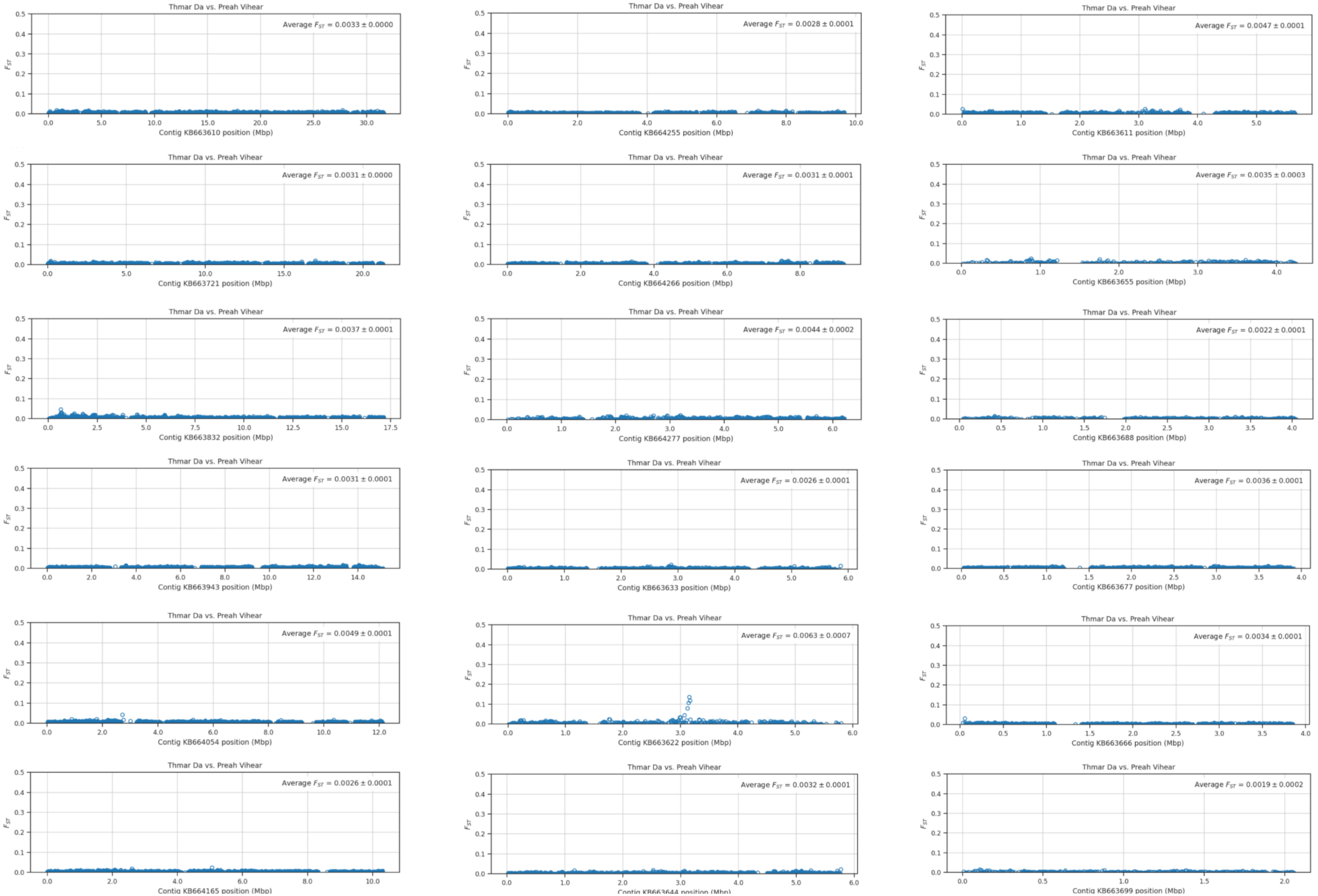

Supplementary Figure 4.2

**TD vs. RK1**

Fst scans in 1000 SNP windows across the largest 18 AminM1 contigs

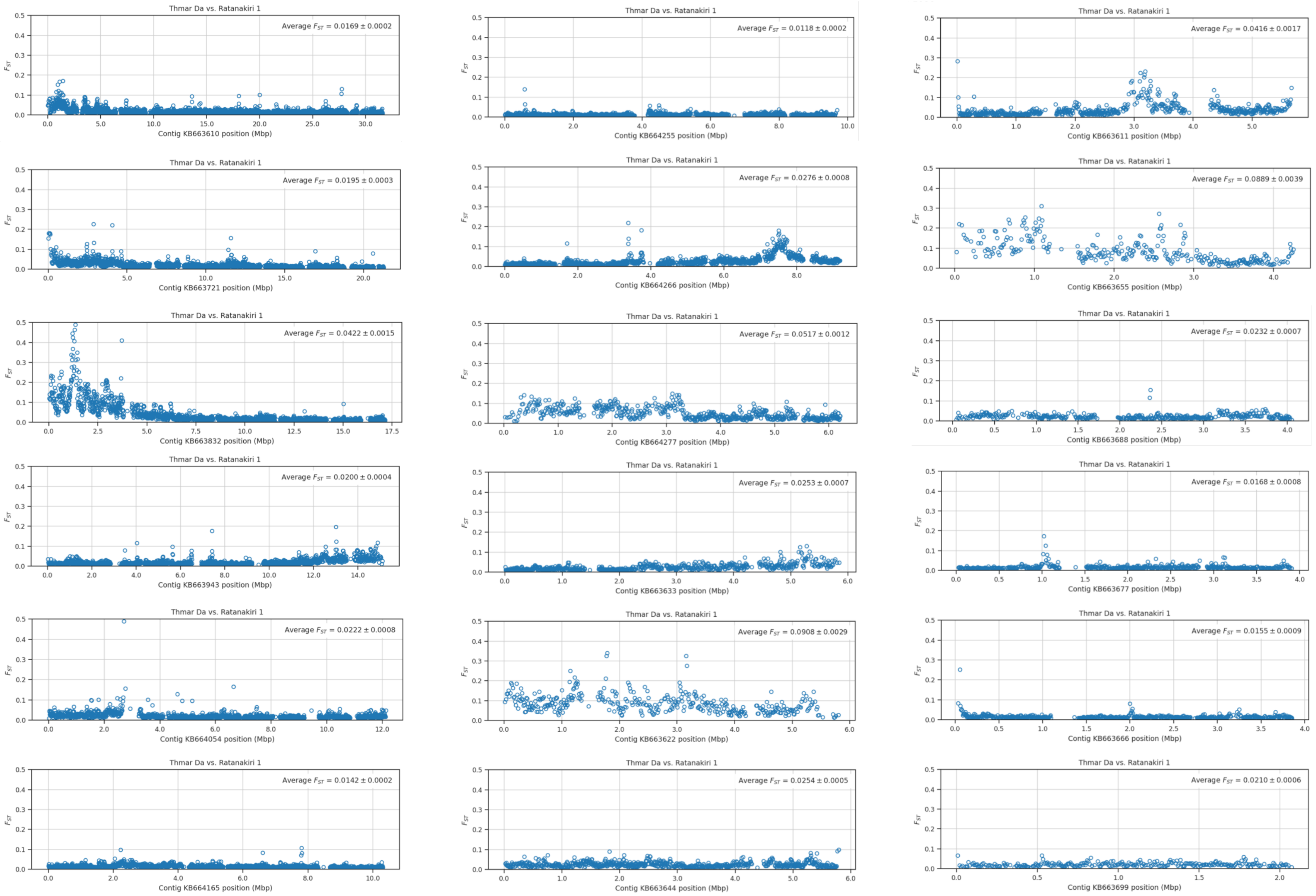

Supplementary Figure 4.3

# TD vs. RK2

Fst scans in 1000 SNP windows across the largest 18 AminM1 contigs

*axes adjusted to accommodate Fst greater than 0.5

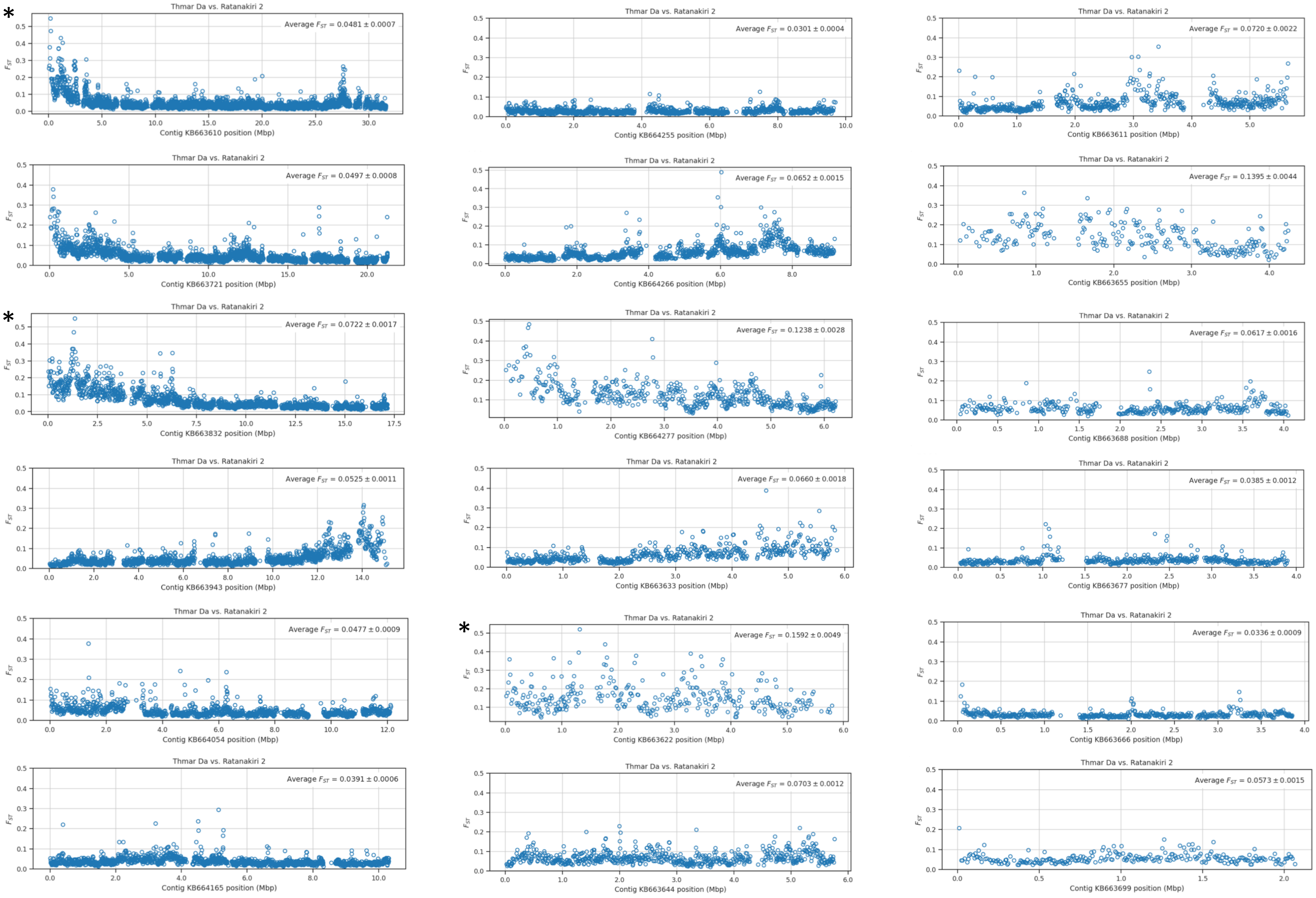

Supplementary Figure 4.4

**PV vs. RK1**

Fst scans in 1000 SNP windows across the largest 18 AminM1 contigs

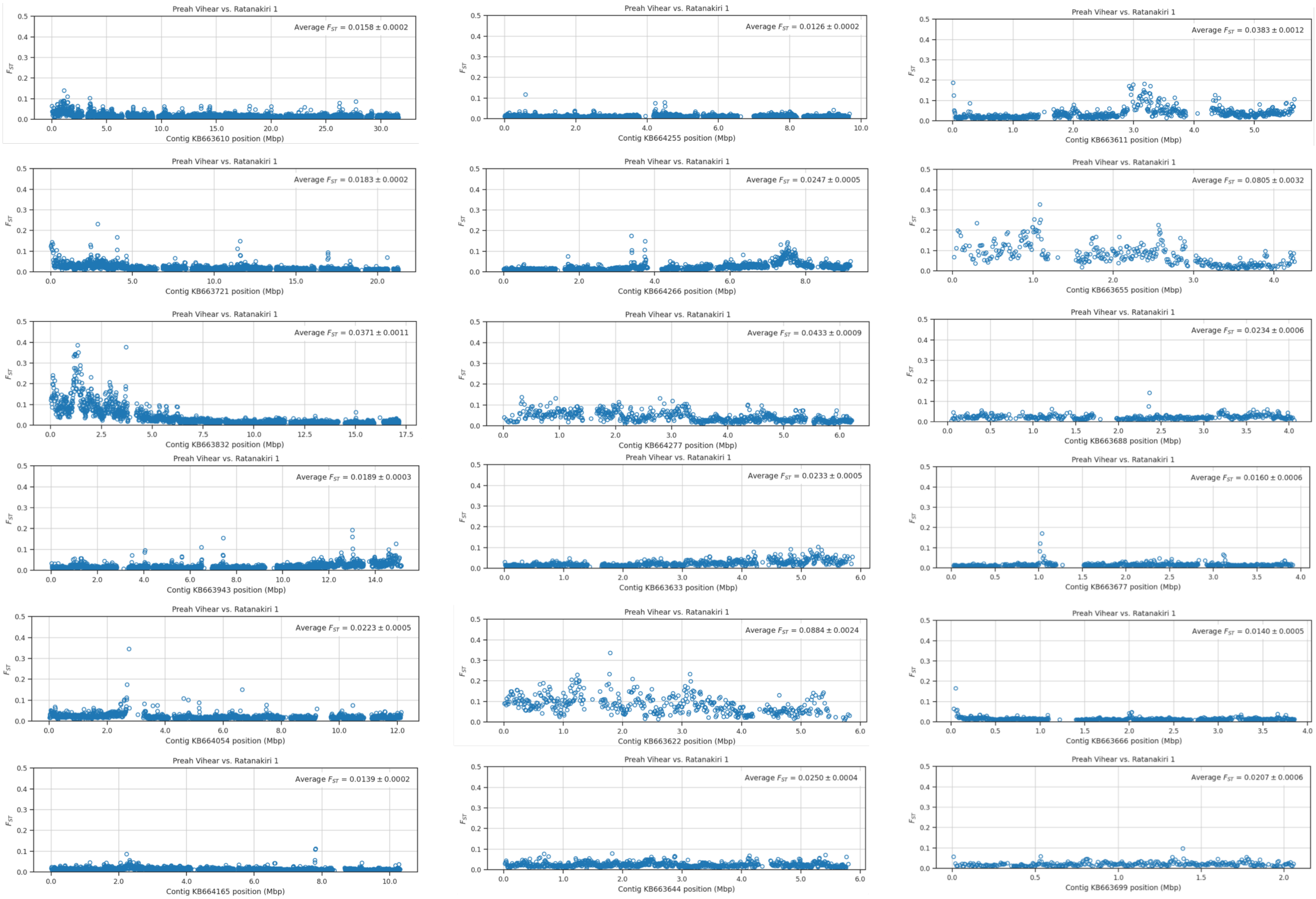

Supplementary Figure 4.5

# PV vs. RK2

Fst scans in 1000 SNP windows across the largest 18 AminM1 contigs

*axes adjusted to accommodate Fst greater than 0.5

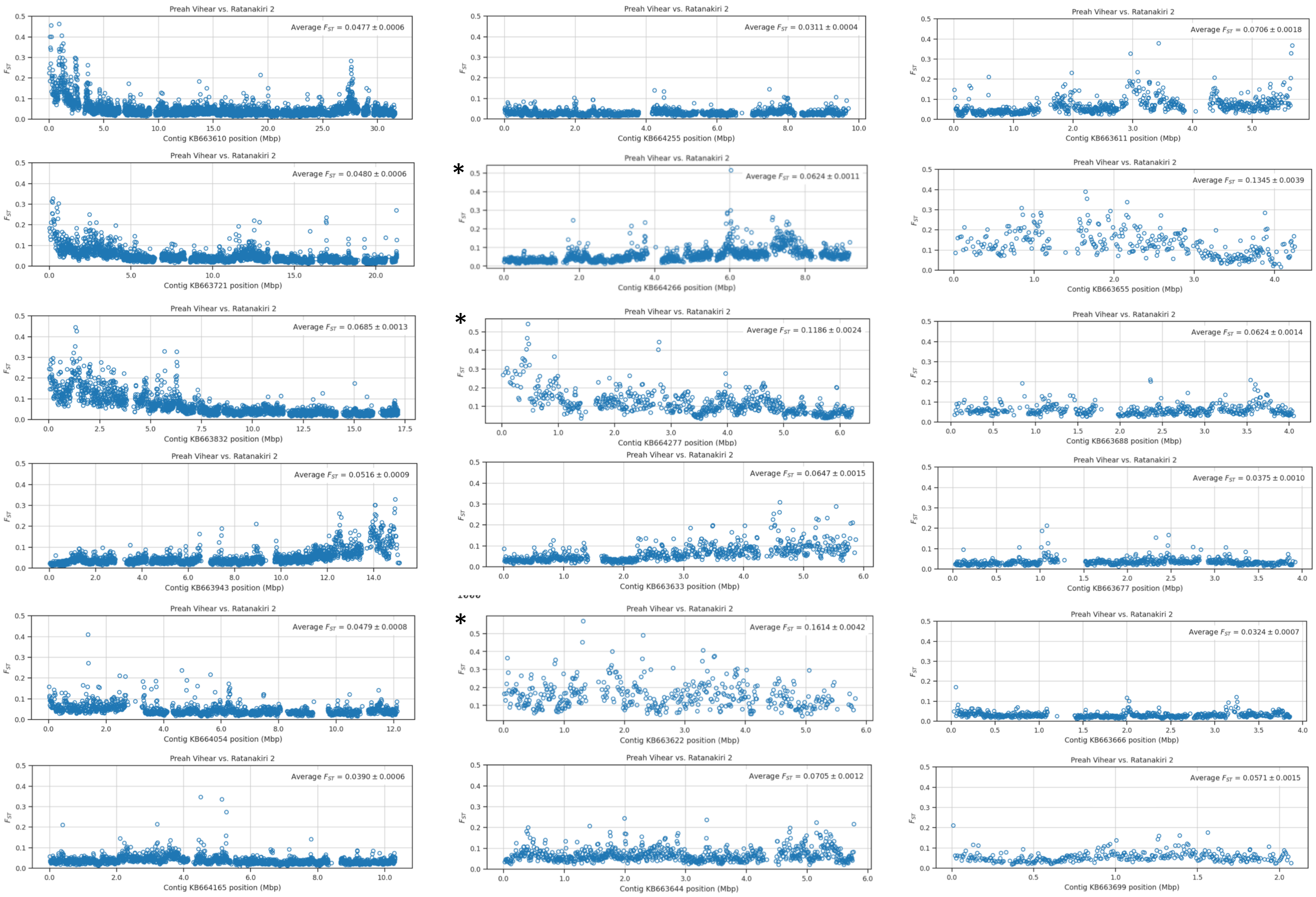

Supplementary Figure 4.6

# RK1 vs. RK2

Fst scans in 1000 SNP windows across the largest 18 AminM1 contigs

*axes adjusted to accommodate Fst greater than 0.5

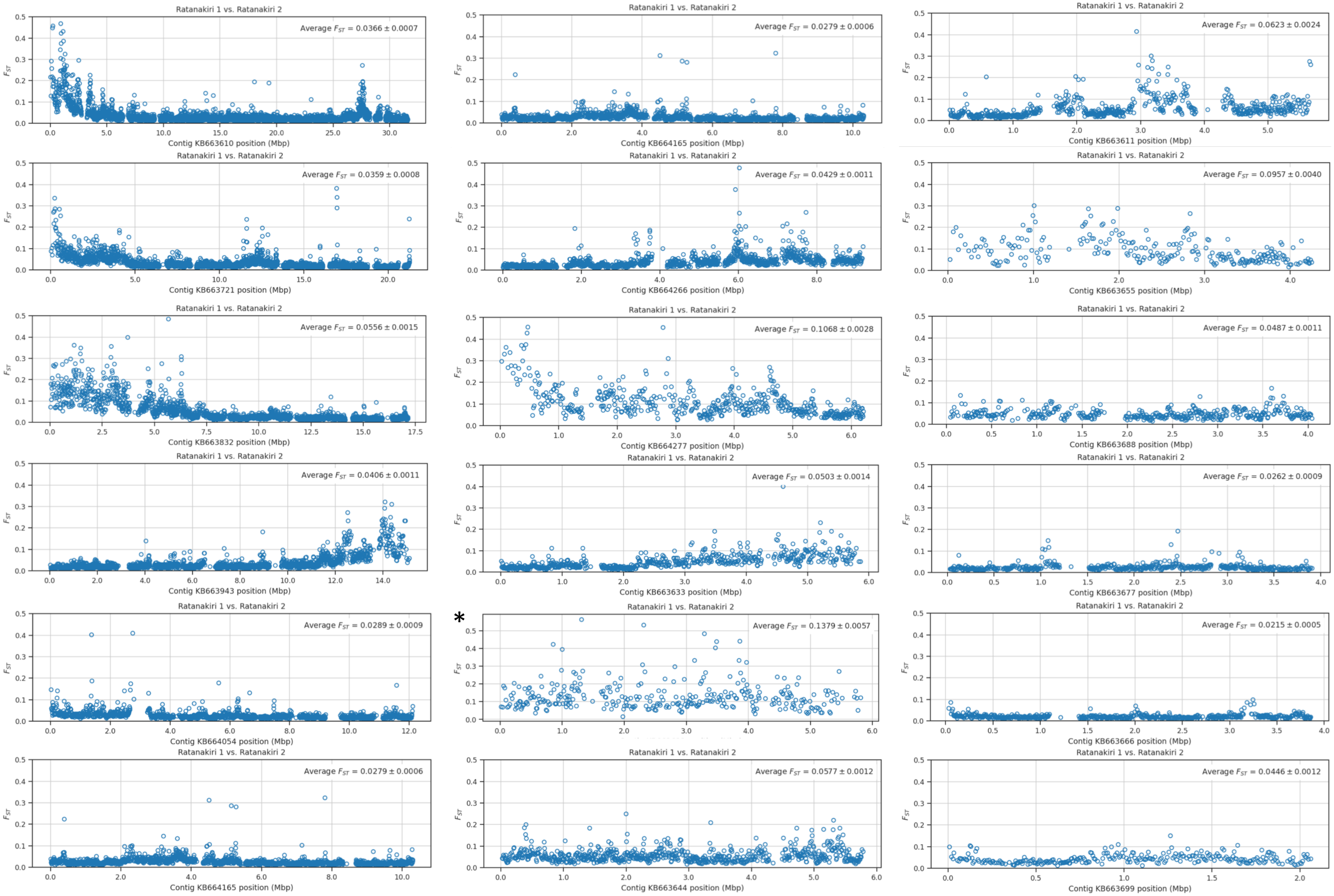
